# Identification of targetable vulnerabilities of PLK1-overexpressing cancers by synthetic dosage lethality

**DOI:** 10.1101/2024.07.18.603978

**Authors:** Chelsea E. Cunningham, Frederick S. Vizeacoumar, Yue Zhang, Liliia Kyrylenko, Peng Gao, Vincent Maranda, He Dong, Jared DW Price, Ashtalakshmi Ganapathysamy, Rithik Hari, Connor Denomy, Simon Both, Konrad Wagner, Yingwen Wu, Faizaan Khan, Shayla Mosley, Angie Chen, Tetiana Katrii, Ben G. E. Zoller, Karthic Rajamanickam, Prachi Walke, Lihui Gong, Hardikkumar Patel, Mary Lazell-Wright, Alain Morejon Morales, Kalpana K. Bhanumathy, Hussain Elhasasna, Renuka Dahiya, Omar Abuhussein, Anton Dmitriev, Tanya Freywald, Erika Prando Munhoz, Anand Krishnan, Eytan Ruppin, Joo Sang Lee, Katharina Rox, Behzad Toosi, Martin Koebel, Mary Kinloch, Laura Hopkins, Cheng Han Lee, Raju Datla, Sunil Yadav, Yuliang Wu, Kristi Baker, Martin Empting, Alexandra K. Kiemer, Andrew Freywald, Franco J. Vizeacoumar

## Abstract

Tumor heterogeneity poses a significant challenge in combating treatment resistance. Despite Polo-like kinase 1 (PLK1) being universally overexpressed in cancers and contributing to chromosomal instability (CIN), direct PLK1 inhibition hasn’t yielded clinical progress. To address this, we utilized the synthetic dosage lethality (SDL) approach, targeting PLK1’s genetic interactions for selective killing of overexpressed tumor cells while mitigating heterogeneity-associated challenges. Employing computational methods, we conducted a genome-wide shRNA screen, identifying 105 SDL candidates. Further in vivo CRISPR screening in a breast cancer xenograft model and in vitro CRISPR analysis validated these candidates. Employing Perturb-seq revealed IGF2BP2/IMP2 as a key SDL hit eliminating PLK1-overexpressing cells. Suppression of IGF2BP2, genetically or pharmacologically, downregulated PLK1 and limited tumor growth. Our findings strongly propose targeting PLK1’s genetic interactions as a promising therapeutic approach, holding broad implications across multiple cancers where PLK1 is overexpressed.

## Introduction

Tumor heterogeneity is an enormous clinical challenge because it provides selective evolutionary advantages to cancer cell subsets, leading to the establishment of aggressive clones that are metastatic and resistant to treatment (*1–3*). Current drug development programmes focus on co-targeting multiple pathways within cancer cells. For example, clonal heterogeneity in colorectal cancer was overcome by simultaneous co-inhibition of MEK and EGFR kinases (*4*). A promising approach to overcome intratumor heterogeneity is based on targeting factors that directly contribute to genetic diversity and intratumor heterogeneity (*5*). This approach should limit the acquisition of multi-drug resistance and treatment failure.

Chromosomal instability (CIN) is one of the key driving forces of genetic diversity within tumors and remains a key underlying feature of genetically diverse malignancies (*6–13*). CIN arises due to aberrant mitotic division, defective double-strand break repair, replication stress or ineffective telomere maintenance (*7, 8, 10, 12, 14–19*). Polo-like kinase 1 (PLK1) is a serine/threonine protein kinase and a central player in controlling CIN (*20–22*). On the molecular level, PLK1 contributes to genome stability by signaling the initiation of mitosis, centrosome maturation, bipolar spindle formation, chromosome segregation, and cytokinesis (*20, 23–26*). Constitutive overexpression of PLK1 leads to CIN and aneuploidy, which are common salient features of most cancers (*22, 27*). In fact, tumor cells upregulate genes such as PLK1 to support their survival and propagation (*28–31*). Previously, we showed that changes in the expression patterns of genes such as PLK1, involved in the maintenance of CIN and factors that remodel the tumor microenvironment, represent some of the earliest events in tumor evolution (*32*). Consistent with this idea, PLK1 is overexpressed in a wide range of cancers, including breast (*33*), colon (*34*), pancreatic (*35*), glioma (*36*), lung (*37*) and prostate (*38*) cancers. However, even after 30 years since its discovery (*39*) and 20 years after its pro-malignant properties were recognized (*40*), bringing PLK1 inhibitors into clinical application has been extremely challenging. The lack of effectiveness of PLK1 inhibitors is due to the difficulty in achieving PLK1-specific inhibition (*24*). Unwanted inhibition of closely related members of the polo-like family can lead to toxicity in the nervous system (*41*) or may interfere with the hypoxic response and promote angiogenesis (*42*). Additionally, PLK1 itself plays key roles in controlling a multitude of cellular processes regulating CIN. Complete loss of function of this protein could be detrimental to normal cells (*24, 43*) and may overcome the tumor-selective basis of treatment. Considering the diverse functions of PLK1 and its family members, there are still major challenges in using direct PLK1 inhibitors in cancer therapy, and alternative strategies are needed.

To take advantage of frequent PLK1 overexpression in tumors, we applied the genetic approach known as synthetic dosage lethality (SDL), in which the overexpression of a gene, such as PLK1, is lethal only when another normally non-lethal genetic alteration is also present (*44, 45*). SDL is still a largely untapped area in cancer research. Since tumor cells upregulate genes such as PLK1, identifying SDL interactions is valuable for revealing new therapeutic targets for cancer treatment (*46–48*). Unlike direct inhibition of PLK1, in which cancer cells may undergo cell death, cell cycle arrest or increased aneuploidy, suppression of molecules that exhibit SDL interactions with PLK1 should result in only cell death **(Figure 1A)**. Here, we report the integration of multiple unbiased platforms, including genome-wide pooled shRNA screening with subsequent validation of SDL targets using pooled *in vivo* and arrayed *in vitro* CRISPR/Cas9 screens in a patient-derived xenograft (PDX) model. As PLK1 overexpression leads to cellular heterogeneity, it is also necessary to confirm whether suppression of its SDL partner(s) truly eliminates PLK1-overexpressing cells. Therefore, we evaluated the effect of the top SDL targets at the single-cell level using a single-cell CRISPR/Cas9 screen (Perturb-seq) with direct capture of guide RNAs (*49*). This work identified IGF2BP2 as a potential new therapeutic target and a novel regulator of PLK1 expression. Subsequent studies were performed using a newly characterized pharmacological inhibitor of IGF2BP2, which was found to affect the expression of PLK1 and preferentially eliminate PLK1-overexpressing cells and tumors.

**Figure 1:**
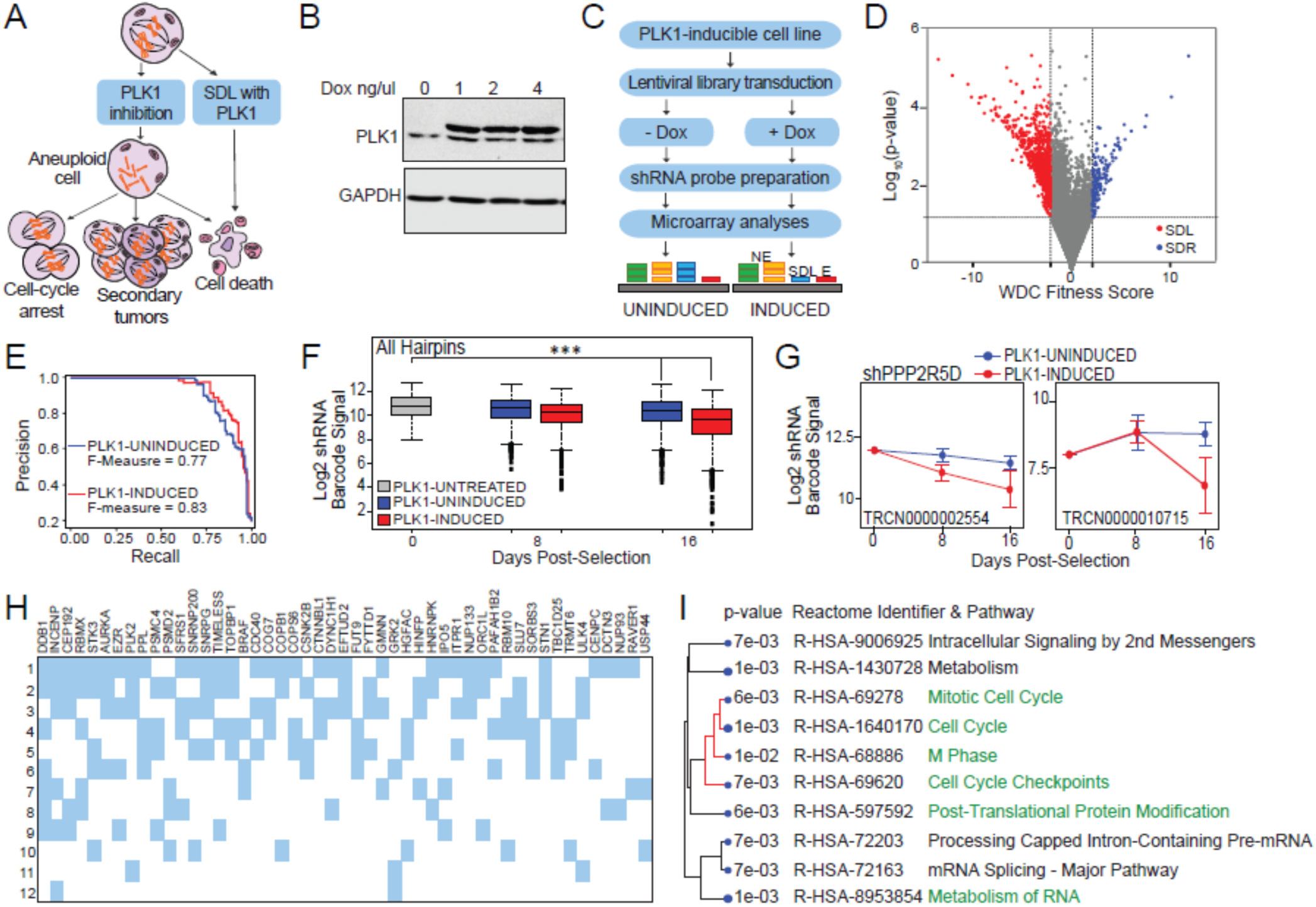
Genome-wide shRNA screening identifies synthetic dosage lethal partners of PLK1. **A.** Schematic illustrating the expected outcome in PLK1-overexpressing cancer cells after inhibition of PLK1 or of the PLK1 SDL target. PLK1 inhibition is expected to cause aneuploidy and potentially an increase in cell death. PLK1-SDL gene inhibition is expected to cause only cell death. **B.** Western blot analysis of the HCT116-PLK1 inducible cell line for PLK1 with and without induction with doxycycline. The upper band represents the constitutively phosphorylated form of the PLK1-S137D mutant. GAPDH was used as a total protein loading control. **C.** Schematic illustration of the genome-wide screening workflow. Example microarray signal outcomes for non-essential (NE, green and yellow), synthetic dosage lethal (SDL, blue), and essential (E, red) genes are shown. **D.** Volcano plot representing the results of the genome-wide pooled shRNA screen. Negative genetic interactions or genes that significantly (p<0.05 and weighted correlation coefficient (WDC) < 2-fold) decreased the fitness scores in the PLK1-overexpressing population compared with the PLK1-nonoverexpressing population are indicated in red. Positive genetic interactions are shown in blue. The total number significant hits came to 960 genes and the full list is provided in Table S1. **E.** Precision vs. recall (PR) curve calculated by measuring the Bayes factor for the genes previously described as general essential and non-essential genes from published screens. F-measure > 0.75. **F.** Box plots summarizing microarray signals for all queried hairpin barcodes at different timepoints in PLK1-untreated (no induction) and PLK1-induced conditions. A 2.2-fold decrease in the induced population from T0 to T16 was observed (Kolmogorov‒Smirnov test p<0.0001). **G.** Drop plots of the microarray signals for the two individual hairpins targeting the PPP2R5D gene over time in the PLK1-uninduced and PLK1-induced strains. **H.** SDL hits that were also picked up from published mitosis-related screens. Only the 45 genes with the most overlap are shown. Rows and columns are sorted by the total number of overlaps in descending order. The references for each row are as follows: 1:PMID: 20360068; 2:PMID: 15616564; 3:PMID: 20360735; 4, 5, 6:PMID: 24104479 in MUS81; BLM; and PTTG1 null cells; 7:PMID: 27929715 in U2OS cells; 8:PMID: 14654843; 9:PMID: 27929715 in RPE1-hTERT cells; 10:PMID: 24104479 in PTEN null cells; 11:PMID: 16564017; and 12:PMID: 17001007 **I**. Dendrograms of the Reactome pathways that are significantly enriched for the 960 PLK1-SDL candidate genes. FDR-adjusted p values are indicated in the figure.

## Results

### Unbiased genome-wide screening revealed novel SDL interactions of polo-like kinase 1

CIN is a hallmark of cancer (*64*), and PLK1 overexpression has been shown to induce this response (*21, 22*). To identify the SDL interactions of PLK1 and exploit them for cancer therapeutics, we selected an inducible system (*65*) with the active PLK1-S137D mutant form in HCT116 cells, a near-diploid model commonly used to study genome instability (*14, 27, 66–72*), as this system does not inherently exhibit any CIN. Unfortunately, established breast cancer cell lines (e.g., MCF-7 and MDA-MB-231) are not ideal for studying genome instability-related events typically associated with PLK1 overexpression due to their extensive aneuploidy. Moreover, the S137D mutant form of PLK1 was previously shown to be defective at the spindle assembly checkpoint (*73*), and checkpoint defects have been linked to CIN (*14*). Thus, as HCT116 cells harbor an intact DNA damage checkpoint and spindle assembly checkpoint (*72, 74*), these cells could serve as an excellent model system for inducing genome instability (because of PLK1 overexpression in our study) and subsequently identifying related SDL interactions **(Figure 1B).**

To identify the gene knockdowns that cause lethality only when PLK1 is overexpressed, HCT116-PLK1 cells were transduced with a pooled lentiviral library of 90,000 shRNA sequences targeting ∼18,000 different genes with ∼5 independent hairpins per gene. Library transduction was performed at a scale of ∼300-fold representation in two distinct populations (induced vs uninduced) as previously described (*50*) **(Figure 1C),** and 960 gene hits were identified **(Figure 1D)** (with at least a 2-fold decrease and p<0.05). The complete list of 960 significant hits, their corresponding weighted differential cumulative change (WDC) scores, and their biological functions are detailed in **Table S1,** and these genes are hereafter referred to as ‘PLK1-SDL’ hits. Although the replicates of the screen showed a high correlation (Pearson correlation coefficient, r>0.9 between replicates) **(Figure S1A),** we compared our screening results to sets of essential and non-essential genes using a previously published framework to ensure that our screening reliably identified true SDL hits (*75*). This approach measured good performance by calculating the accuracy score from the precision and recall test (F-measures > 0.75) **(Figure 1E)**. The cumulative signal used to calculate the fitness score of all PLK1-SDL gene hits displayed a significant 2.2-fold decrease in the induced population from T0 to T16 (Kolmogorov–Smirnov test p<0.0001) as opposed to the uninduced population from T0 to T16, which had a 1.3-fold decrease (Kolmogorov–Smirnov test, p<0.0001) **(Figure 1F)**. Consistent with this, the overlay of all individual signals from all the dropouts was highly represented in the induced population compared to the uninduced population (**Figure S1B)**. To illustrate the dropout of signals over time in the induced population alone, we analyzed one of the top hits, PPP2R5D, a regulatory subunit of the protein phosphatase 2A (PP2A) complex, which we previously reported to exhibit SDL with PLK1 (*27*) **(Figure 1G)**, reiterating the confidence in our screening approach. SDL interactions are functionally coherent (*44*), and as expected, upon comparison of our PLK1-SDL hits with previously published mitosis-related screens (*76–83*), we found several hits associated with mitosis, DNA repair, and cell cycle-related pathways, in addition to several completely novel PLK1 partners **(Figure 1H and Table S2).** Consistent with these findings, Reactome pathway analyses indicated that PLK-SDL hits were enriched in the cell cycle (adj. p 1e-03), mitosis- and checkpoint-related pathways (adj. p < 6e-03), and RNA metabolism pathway (adj. p 1e-03) **(Figure 1I)**. To gain insight into the functional relevance of the SDL hits, thorough literature analyses of the 960 genes were performed using Cytoscape Search Tool for the Retrieval of Lengths (STRING) analyses. While these studies categorized the SDL hits into multiple biological functions, including cell cycle progression, centrosome amplification and cytokinetic component enrichment, the extensive interactions among these components indicated that most of the SDL interactions were also functionally related **(Figure S1C; Table S1)**.

While the enrichment analyses increased confidence in our findings, we also used drug response data from the Genomic of Drug Sensitivity in Cancer (GDSC) (https://www.cancerrxgene.org), as we found that several of the SDL hits had chemical inhibitors. This analysis revealed that a few inhibitors of SDL hits selectively suppressing PLK1 overexpressing cells **(Figure S2A)**. Thus, our screen revealed that GSK3A and its inhibitor CHIR-99021 preferentially affect PLK1-overexpressing cells. Similarly, ALK receptor tyrosine kinase was found among the SDL hits, and the ALK inhibitor NVP-TAE684 appeared to selectively target PLK1-overexpressing cells **(Figure S2A).** Given that ALK inhibition activates the spindle assembly checkpoint, causing mitotic delay (*84*), and that PLK1-overexpressing cells may be defective in the spindle assembly checkpoint (*73*), we speculate that the inhibition of ALK in PLK1-overexpressing cells may lead to mitotic catastrophe. These completely independent cross-validations by chemical genetics and testable hypotheses confirm the quality of our screens and their potential to identify new potentially therapeutically relevant targets.

### Systematic prioritization of PLK1-SDL hits for further validation studies

Genome-wide screens tend to produce some false positive hits that can confound SDL identity. Most genome-wide studies cherry-pick one or two hits to serve as validated proofs of principle, but most of the data often remain unvalidated and therefore underutilized by the research community. However, validating all hits from a large-scale screen, such as the one conducted here and pointing toward 960 SDL hits, is also practically challenging. Therefore, we applied three distinct strategies that allowed us to prioritize a subset of PLK1-SDL hits for experimental validation while also maintaining the strength of a large-scale unbiased screening approach.

Our first strategy to prioritize PLK1-SDL hits was based on the rationale that SDL genes that are differentially upregulated when PLK1 is overexpressed represent coregulatory mechanisms that become essential within the molecular context of elevated PLK1 levels (*85*). Therefore, we used patient data from The Cancer Genome Atlas (TCGA) (https://portal.gdc.cancer.gov/) to determine the number of PLK1-SDL hits that were co-upregulated with PLK1 across 33 different cancer types. We found that the expression of a subset of PLK1-SDL hits was positively correlated with PLK1 levels across several cancer types, suggesting that these hits may represent specific genetic dependencies of PLK1-overexpressing cells **(Figure 2A)**. After filtering out essential genes (*56, 86*) and genes that are unlikely to be expressed in HCT116 cells (based on CCLE data (*54*); log2 expression score <5.0), we selected 20 genes whose expression strongly correlated with PLK1 levels (Spearman-rank correlation coefficient, r ≥ 0.4 in a majority of cancer types) **(Figure 2A).** In our second strategy, we investigated how many of the 960 PLK1 SDL hits were ‘pre-validated’ in independently published essentiality screens (*55, 56*). All the cell lines from these published studies were classified into two groups, those with high PLK1 expression (top 10%) and those with low PLK1 expression (bottom 10%), after which we determined whether our PLK1-SDL hits were more essential in the naturally PLK1-overexpressing cell line group. We found 30 SDL hits in the Marcotte *et al*. data **(Figure 2B)** and 19 hits in the Project Achilles data **(Figure 2C)** that had higher essentiality scores in PLK1-overexpressing cells (Wilcoxon rank sum test, p<0.05) (*55, 56*). The complete list with p values is provided in **Table S3.**

**Figure 2:**
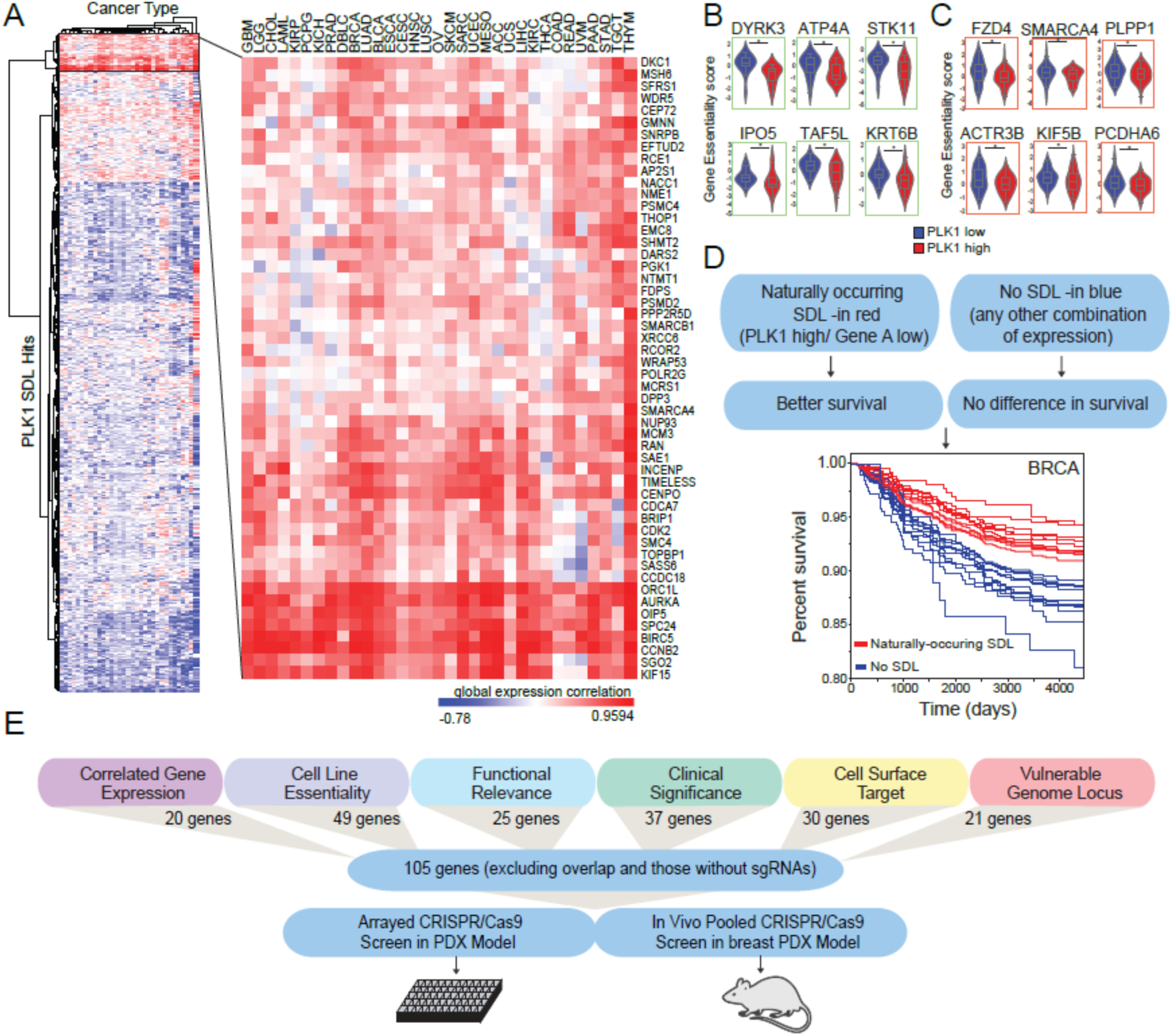
Prioritization of the PLK1-SDL candidate genes using computational analyses. **A.** Hierarchical clustering of the correlation between the expression of PLK1 and the expression of each SDL hit across 33 cancer types from The Cancer Genome Atlas (TCGA) patient data obtained from Genomic Data Commons (GDC). **B and C.** Violin plots of the difference in essentiality scores for PLK1 SDL hits in different cell lines grouped by low versus high PLK1 expression. Essentiality scores derived from published shRNA screening data (Marcotte *et al*., highlighted in green) and the Project Achilles database (https://depmap.org/portal/achilles/) (highlighted in orange). The p value significance was calculated using the non-parametric Wilcoxon rank sum test. **D.** Schematic of the identification of naturally occurring SDL interactions in patients with breast cancer (BRCA) using Kaplan‒Meier survival curves. Log rank p values were computed to calculate significance. **E.** Schematic summarizing the gene prioritization procedure for experimental validation.

Our third approach assessed SDL interactions based on patient prognoses associated with the expression level of the screen hits (*85*). Briefly, patient samples were queried for ‘naturally occurring SDL interactions’, depending on the expression levels of PLK1 and its SDL partner **(Figure 2D).** The genes for which naturally occurring SDL expression patterns were positively correlated with significantly improved patient survival (37 genes; Kaplan‒Meier log-rank test, p value <0.05) were selected for further validation **(Figure 2D and S2B; Table S4)**. Apart from the above three approaches, we also included 21 genes from the short arm of chromosome 19 (19p13.2-3), as our screen identified many genes from this locus **(Table S5)**. Given that this region has been associated with macrocephaly (*87*) and that PLK1 and its centrosome functions have been previously linked to both micro- and macrocephaly (*88, 89*), we expected to see functional crosstalk between PLK1 and this vulnerable locus. We also included 25 functionally relevant PLK1-SDL hits, as determined through gene prioritization algorithms (*57*) **(Table S5).** Finally, 30 potential cell surface SDL targets were included since they may represent targetable vulnerabilities for advanced strategies such as antibody-based inhibitors **(Table S5)**. Overall, after removing hits overlapping among these approaches, we selected 134 genes for further studies. Since 29 of these genes failed to guide RNA cloning, we focused our efforts on validating the remaining 105 genes **(Figure 2E).**

### Combination of *in vivo* pooled CRISPR and *in vitro* arrayed CRISPR screens in a PLK1-overexpressing breast cancer PDX model

Following the prioritization of 105 candidate SDL hits, we made a systematic effort to identify the best SDL hits as potential antitumor targets. To this end, we selected 210 sgRNA sequences for 105 genes and queried the essentiality of the prioritized PLK1-SDL hits in a previously described breast cancer PDX model, HCI-010, which closely mimics tumor properties (*58*). The derived cells displayed PLK1 overexpression compared to a non-malignant breast cell line Hs578Bst **(Figure 3A).** We engineered HCI-010 cells to stably express Cas9 and used them in a CRISPR-based arrayed screen following lentiviral transduction and high-throughput imaging **(Figure 3B)**. To evaluate each SDL hit, HCI-010 cells with or without Cas9 expression **(Figure 3B)** were transduced with pooled lentiviral particles expressing two independent sgRNAs in a single well. The cells were monitored over time to determine the viability of PLK1-overexpressing HCI-010 cells after gene knockout by automated image analysis **(Figure 3C)**. The confluency of each well was calculated using Molecular Devices MetaXpress image analysis software and used as a proxy measurement for cell viability. In total, 60 of the 105 genes were associated with an ∼40% or greater reduction in confluency (Student’s t test, p<0.0.5) **(Figure 3D and S3A; Table S6)**.

**Figure 3:**
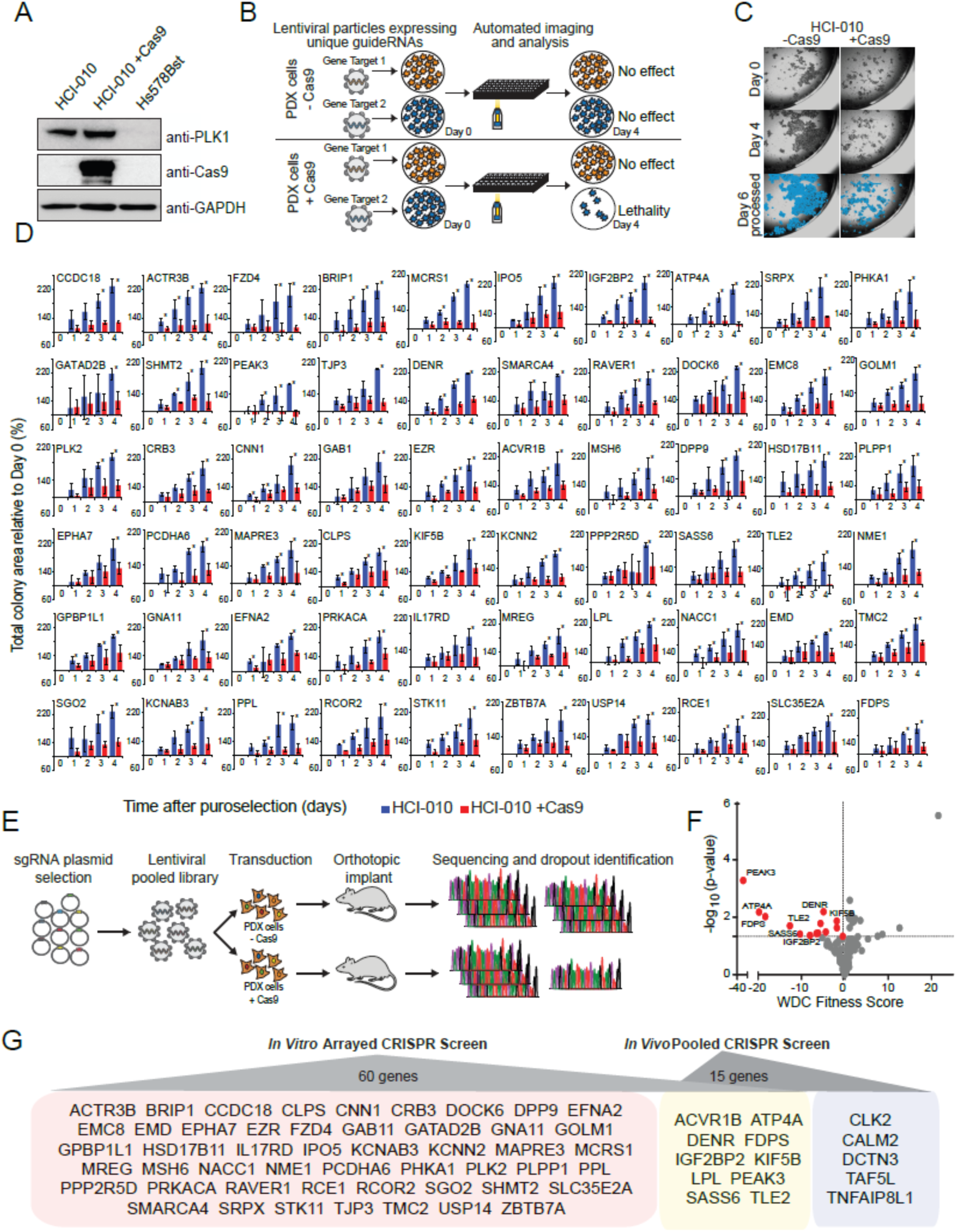
CRISPR-based knockout validation of PLK1-SDL candidate genes in a patient-derived breast cancer model. **A.** Western blot analysis of PLK1 and Cas9 in HCI-010 cells, HCI-010 cells stably transduced with a Cas9 expression vector, and Hs578Bst non-malignant breast epithelial cells. GAPDH is shown as the total protein loading control. **B.** Schematic of the methodology for CRISPR arrayed *in vitro* screening validation in PDX breast cancer cells with and without Cas9 using high-throughput imaging automation to determine lethality. **C.** Sample images acquired using automated imaging over time and analysis of different PLK1-SDL candidate gene sgRNAs in -Cas9 and +Cas9 HCI-010 cells. The MetaXpress object masking overlay is shown in blue for the day 6 images. **D.** Bar graph quantification of the imaging analysis over the course of 4 days. n = 3, * p value < 0.05, Student’s t test. **E.** Schematic of the *in vivo* pooled CRISPR screening in a PDX breast cancer model with and without Cas9. Sequencing was performed to capture *sg*RNA dropout. **F.** Volcano plot of the log10 p value versus the weighted correlation coefficient (WDC) fitness score for all queried genes from the *in vivo* pooled CRISPR screen. The P value cutoff was < 0.05 based on WDC permutation shuffling. Significant essential genes are colored red. **G.** The final list of SDL hits identified from the *in vitro* arrayed CRISPR screen and *in vivo* pooled CRISPR screen.

In parallel, we generated a pooled lentiviral library containing all 210 sgRNA sequences and transduced HCI-010 PDX cells with or without Cas9 expression in a manner mimicking the genome-wide pooled screen, as we described previously (*90*) **(Figure 3E)**. Following puromycin selection, the sgRNA-transduced HCI-010 cells were introduced into the mammary fat pad regions of female mice with more than 4000-fold representation per sgRNA (3 million cells injected containing 210 library sgRNAs). After the tumors had grown for three weeks, they were harvested, and the genomic DNA was extracted. The sgRNA sequences were PCR amplified with Illumina adapter sequences and sequenced to determine the dropout of sgRNAs from the HCI-010 Cas9-expressing samples, while HCI-010 tumors without Cas9 expression served as a baseline control. Replicates of the screen showed a high correlation (Pearson correlation coefficient, r > 0.9) **(Figure S3B)** between seven tumors in each condition. This *in vivo* pooled CRISPR screen identified 15 SDL hits (p<0.05) **(Figure 3F and Table S7)**. Overall, by these two complementing approaches, we validated 65 PLK1-SDL hits in the HCI-010 PDX model, with 10 SDL hits overlapping in *in vivo* and *in vitro* experiments **(Figure 3G)**. Cas9 genome editing was confirmed in these experiments to eliminate off-target effects by using a cleavage detection assay **(Figure S3C)**. While these 65 SDL hits were found to be essential for the survival of PLK1-overexpressing cells, at this stage, we did not test them for their essentiality in PLK1^low^ non-malignant controls. As the PLK1^low^ non-malignant breast cell line Hs578Bst grows very slowly, we decided to test only the best SDL hits after further filtering them.

### Shortlisting of the top candidates via Perturb-seq with direct guide RNA capture

PLK1 overexpression is known to induce CIN and heterogeneity in tumor cells (*21, 22*). Therefore, we next asked which of the 65 SDL hits eliminated most of the PLK1-overexpressing single cells. The direct-capture Perturb-seq methodology queries the changes within the transcriptome at the single-cell level (*49*) and can also be used to evaluate the survival of individual CRISPR knock-outs at the single-cell level, akin to our negative-selection genome-wide screen **(Figure 1C and 1D)**. To take advantage of this approach, we constructed a new sgRNA library targeting 65 experimentally validated genes in the Chromium Single-cell Gene Expression v3.1 (Next GEM)-compatible LV13 vector backbone (U6-gRNA:EF1a-Puro-2a-BFP; MilliporeSigma) with 10x Genomics-compatible capture sequence 1 (CS1: 5’-TTGCTAGGACCGGCCT TAAAGC-3’) at the stem‒loop position. In this system, the functionally expressed sgRNA incorporates a capture sequence directly into the guide scaffold to allow direct capture of sgRNAs during single-cell RNAseq (*49*). Sixteen guide sequences targeting housekeeping genes, such as RPL3, RPL11, RPL13 and RPL18, were used as positive controls, as described previously (*75*), and 15 non-targeting guide sequences were used as negative controls **(Table S8)**. This sub-library of 161 guides targeting 65 genes and the relevant controls were used to transduce innately PLK1-overexpressing MCF7 breast cancer cells (*91*) **(Figure 4A)**.

**Figure 4:**
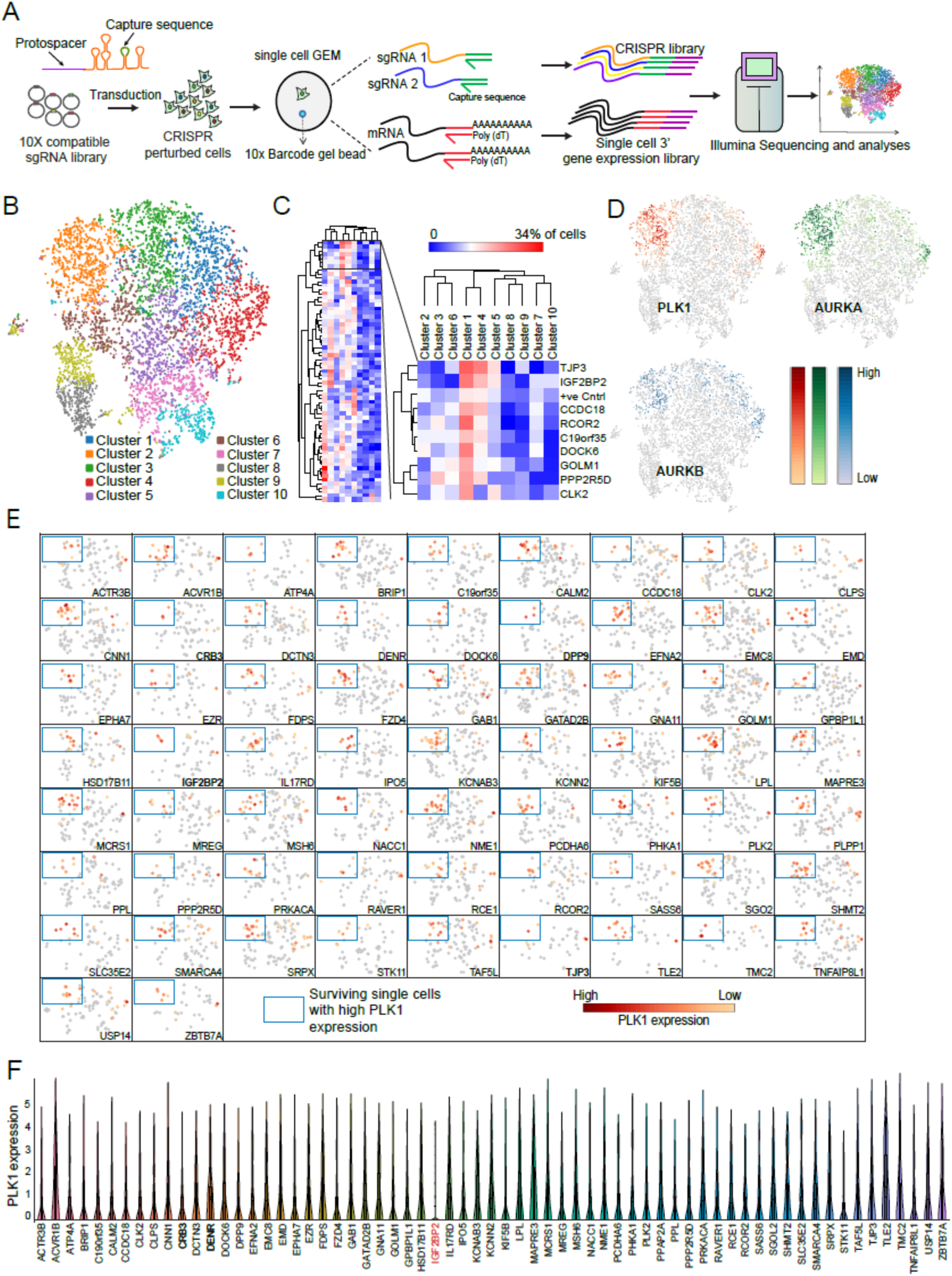
Shortlisting of the top PLK1-SDL candidates using direct guide RNA capture Perturb-seq screening. **A.** Schematic overview of the Perturb-seq screening with direct capture of the guide RNA workflow. Briefly, a 10X compatible guide library with direct capture sequence 1 in the stem loop region was transduced into MCF7 cells, which were subsequently grown for four days following puromycin selection. The gene expression library and CRISPR KO library were prepared following barcoding and indexing of single cells and sequenced with NovaSeq. **B.** Single-cell K-means cluster projection of the T-Sne-embedded pooled sgRNA screen showing 10 clusters (n=7434 cells). **C.** Hierarchical clustering using Pearson correlation analysis of the percentage of knockout cells for each gene in the 10 different clusters. Knockouts of a few genes that are enriched in clusters 1 and 4, along with positive controls, are presented. **D.** Expression analyses of PLK1, AURKA and AURKB in individual cells on a log2 scale (n=7434 cells) show that they are highly co-expressed in cluster 2. **E.** Cells with high PLK1 expression (marking cluster 2 with a box) were tracked for each of the 65 single knockouts. Knockouts of CRB2, DPP9, IGF2BP2, and TJP3 are associated with the lowest numbers of cells with high PLK1 levels (negative selection). **F.** Violin plots showing PLK1 levels in the aggregates of all single cells from 65 knockouts. The graph was generated by 10x Genomics Loupe Browser software.

Following transduction and selection, pooled knockout cells were cultured for four days, and single-cell gel beads in emulsion (GEMs) with barcoded polyadenylated mRNA primers and sgRNA capture sequences were generated for single-cell transcriptome and CRISPR library construction. After sequencing and standard data processing using the 10x Genomics Cell Ranger (ver. 6.1.2) software package, the UMI matrix of cells from each replicate was extracted with single guides, subjected to quality control and normalization, and subsequently imported into the Loupe browser (ver. 6.0) for downstream analysis and t-sne plot generation **(Figure 4B)**. We sequenced 22,041 single cells that yielded an average of 51,479 reads per cell, with a median of ∼4000 genes expressed per cell. Individual cells that had fewer than 5000 transcripts (<5000 UMIs per cell) or were mapped to 0 or ≥ 2 guides in a single cell were eliminated. Overall, we considered 7434 single cells with at least 50 representative cells for most of the knockouts when we mapped a single guide per cell **(Figure S4A).** The efficiency of each knockout was confirmed by comparing the expression of the corresponding knockout gene with that of negative controls **(Figure S4B).** The expression of six genes (LPL, CNN1, C19ORF35, CLPS, ATP4A and TMC) did not significantly decrease, and these genes were excluded from further analyses **(Figure S4B).** The resulting t-SNE plot from the correlation of transcriptomic data across 65 gene knockouts outlined in 10 clusters **(Figure 4B)** with clusters 1 and 4 enriched for knockouts of eight genes (*TJP3, IGF2BP2, CCDC18, RCOR2, DOCK6, GOLM1, PPP2R5D* and *CLK2*), included positive controls **(Figure 4C)**. As the housekeeping positive controls are expected to cause cell death, co-enrichment of these knockouts in clusters 1 and 4 with the positive controls indicated that the loss of functions of these genes aligns with the expected SDL phenotype. Moreover, enrichment of PPP2R5D knockouts within these clusters also increases the confidence in our approach, as we previously reported that PPP2R5D exhibits SDL with PLK1 (*27*). Interestingly, cluster 2 was enriched not only for PLK1-overexpressing cells but also for Aurora kinase-overexpressing cells **(Figure 4D)**. We therefore mapped individual gene knockouts separately to determine their relationship with cell populations with different PLK1 levels **(Figure 4E).** We found knockouts of four genes (*IGF2BP2, CRB3, DPP9 and TJP3*) that displayed negative selection with fewer PLK1-overexpressing cells or increased lethality to cells with higher PLK1 levels **(Figure 4E)**. Overall, these results indicate that Perturb-seq screens with direct guide RNA capture efficiently revealed few potential candidates for further analyses.

### Loss of IGF2BP2 affects PLK1 expression and suppresses PLK1-overexpressing cells, tumorspheres and tumors

As PLK1-overexpressing cancer cells may exhibit high PLK1 activity, we tested whether the loss of IGF2BP2, *CRB3, DPP9 or TJP3* affects PLK1 mRNA levels. CRB3, DPP9 and TJP3 knockout did not affect PLK1 expression, but PLK1 expression was decreased in cells with IGF2BP2 knockout **(Figure 4F)**. To confirm this, we performed digital droplet PCR (ddPCR) to determine the absolute expression levels of both IGF2BP2 and PLK1 after two days of IGF2BP2 knockdown. We found PLK1 expression to be significantly downregulated in response to IGF2BP2 silencing in PLK1-overexpressing MCF7 (p<0.001) and BT549 cells (p<0.05) **(Figures 5A and S5A, S5B and S5C)**. In contrast, PLK1 transcript levels increased in MDA-MB-231 cells (p<0.001) when IGF2BP2 was knocked down **(Figure 5A)**. Thus, it appears that the loss of IGF2BP2 can differentially regulate PLK1 mRNA in distinct biological contexts. Nevertheless, we observed a decrease in PLK1 protein levels **(Figure 5B)** following IGF2BP2 silencing in all the tested cell lines, including MDA-MB-231 cells. To determine whether the functional relationship between PLK1 and IGF2BP2 is restricted to specific cell line models, we monitored PLK1 and IGF2BP2 at the protein level and evaluated the correlation between PLK1 and IGF2BP2 mRNA levels in multiple tumor types using The Cancer Genome Atlas (TCGA) data. We found a strong positive correlation between the expression of these two genes (Spearman-rank correlation coefficient, r ≥ 0.3) in multiple cancers (**Figure 5C)**. This correlation suggested that the SDL interaction between PLK1 and IGF2BP2 is based on a reduction in PLK1 activity in the absence of IGF2BP2, which preferentially affects PLK1-overexpressing cells that become dependent on high PLK1 levels.

**Figure 5:**
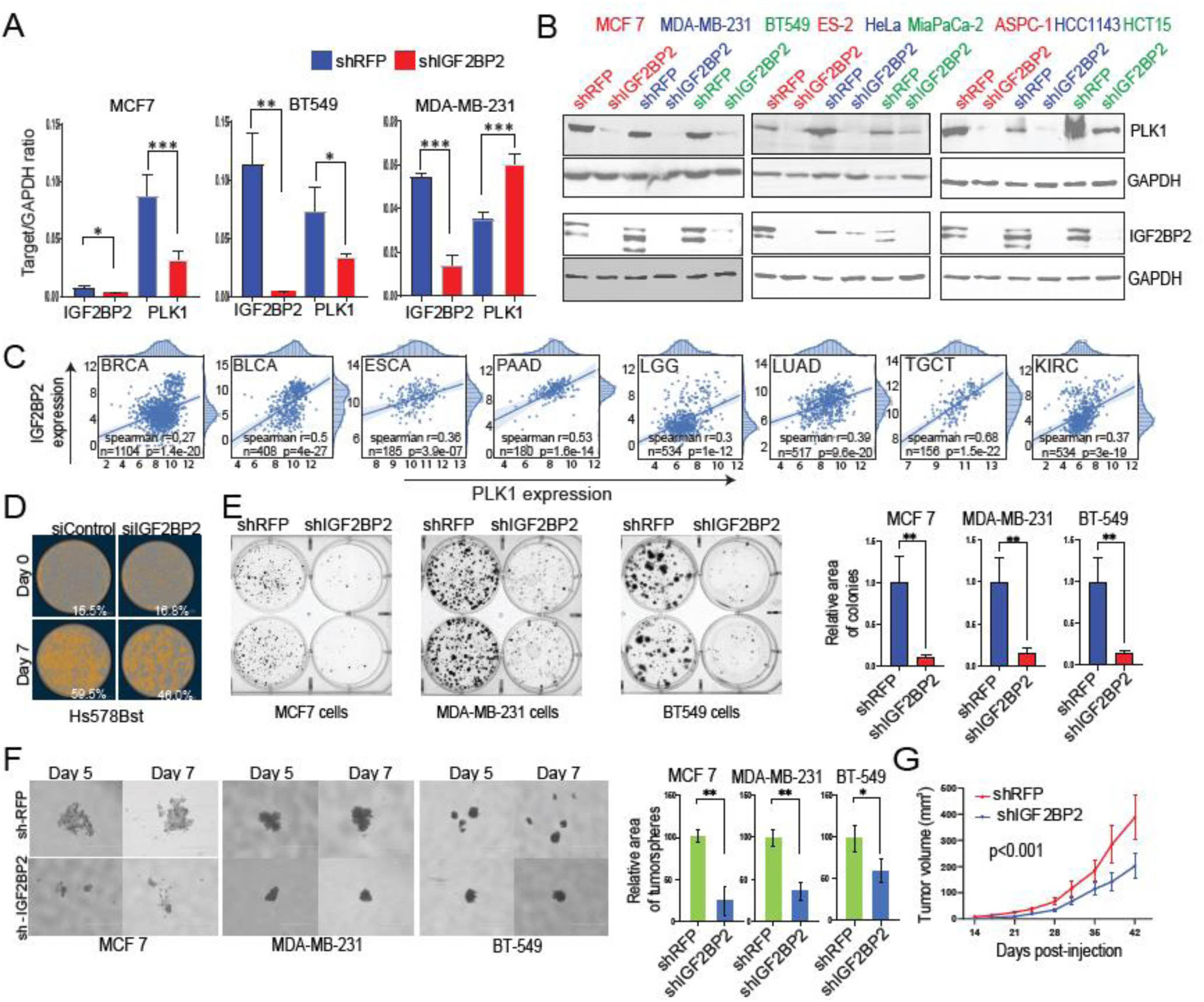
IGF2BP2 affects the expression of PLK1 and tumor growth. **A.** Absolute quantification results of ddPCR are presented as target/GAPDH ratios. After knocking down IGF2BP2, PLK1 expression was significantly downregulated in MCF7 (P<0.001) and BT549 cells (P<0.05). In contrast, PLK1 transcript levels increased in MDA-MB-231 cells (P<0.001). **B.** Western blot showing the levels of PLK1 following the knockdown of IGF2BP2. GAPDH was used as a loading control. **C.** The expression patterns of PLK1 and IGF2BP2 in multiple cancer types from TCGA were positively correlated. Spearman rank correlations were calculated using the log2 scale and the RNA-Seq Expectation Maximization (RSEM)-normalized expression of PLK1 and IGF2BP2. The frequency plots along the x-axis (top) show the frequency of PLK1 expression, and the frequency plots along the y-axis (right) show the frequency of IGF2BP2 expression in each tissue type. Each patient is represented by a blue dot. The Spearman correlation coefficient (r), number of patients (n), and p value significance (p) are included in the graphs. The blue line represents the best linear fit of the distribution. **D.** Sample images with cell masking shown in orange and a box plot of Hs578Bst breast epithelial cell confluency before and after transfection with IGF2BP2-targeting *si*RNA. No significant (n.s.) decrease in confluency was observed relative to that of the non-targeting *si*RNA control, and HS578Bst cells exhibited normal expansion. **E.** Colony formation assay was performed with the indicated breast cancer cells following knockdown of IGF2BP2. shRFP was used as a control. The colonies were quantified using ImageJ software for the area of colonies representing the overall colony abundance compared to the area of the control in MCF7 (p<0.01), MDA-MB-231 (p<0.01), and BT549 (p<0.05) cells. **F.** Representative images of tumorspheres formed from MCF7, MDA-MB-231, and BT549 cells (scale bar, 1000 μm). Images of tumorspheres were taken using an EVOS m5000, and the tumorsphere area was calculated using ImageJ software. The graph represents the area of tumorspheres after knocking down IGF2BP2 compared to that of the corresponding control in MCF7 cells (p<0.01), MDA-MB-231 cells (p<0.01), and BT549 cells (p<0.05). **G.** Graph representing the effect of IGF2BP2 knockdown on tumor volume. MDA-MB-231 cells (2×106) transduced with shIGF2BP2 or shRFP (control) were injected into the mammary fat pads of immunodeficient female NOD-SCID mice (n=10/group), and the tumor volume was measured every 3 to 4 days. The graph shows the mean tumor volume (± standard error) at different time points post-injection. The IGF2BP2 knockdown group exhibited a significant decrease in tumor volume compared to that of the control group (p<0.001, two-way ANOVA).

Importantly, unlike direct PLK1 inhibition, which leads to complete disruption of PLK1 function, thereby affecting normal cells, downregulation of PLK1 by targeting IGF2BP2 still results in minimal PLK1 protein levels, possibly without abrogating its functions in normal cells. To test this possibility, we confirmed that the loss of IGF2BP2 does not affect non-malignant Hs578Bst cells that do not overexpress PLK1 **(Figure 3A and 5D)**. As Hs578Bst cells did not grow efficiently with lentiviral knockdown, we used siRNA to transfect these cells. The cells were transfected with siRNA, and confluency was monitored using live-cell imaging (S3-IncuCyte®). Knockdown of IGF2BP2 did not affect cell growth compared to that of the corresponding control cells when monitored over seven days **(Figure 5D).** Using the same live-cell imaging technique, we also used the HCT116-PLK1 inducible model cell line from the genome-wide screen and found that the loss of IGF2BP2 caused a decrease in viability only in the PLK1-induced condition over a period of 10 days, which matched the results of the genome-wide screen **(Figure S5D).** Furthermore, compared with control shRNA-transduced cells, knockdown of IGF2BP2 also significantly decreased colony formation in a panel of breast cancer cell lines, including MDA-MB-231 cells, which presented an increase in PLK1 mRNA and a decrease in PLK1 protein abundance upon IGF2BP2 knockdown **(Figure 5E and S5E).** Finally, to investigate how IGF2BP2 knockdown affects cancer stem cells, we performed tumorsphere analysis in selected breast cancer cell lines **(Figure 5F)**. Tumorsphere models were used since they better simulate tumor biology than cells cultivated in monolayers, and knockdown of IGF2BP2 effectively suppressed the growth of breast cancer cells in tumorspheres. Following these findings, we next examined whether IGF2BP2 loss reduces tumor growth in xenograft models. To test the ability of IGF2BP2 knockdown to reduce tumor size *in vivo*, we used individual shRNAs to silence IGF2BP2 in MDA-MB-231 cells and injected the cells into immunodeficient female NOD-SCID mice. Consistent with the findings observed in the tumorsphere models, silencing IGF2BP2 reduced the growth of PLK1-overexpressing MDA-MB-231 tumors, suggesting that this model of human TNBC was successful **(Figure 5G).**

### Pharmacological inhibition of IGF2BP2 affects PLK1 expression and decreases the expansion of PLK1-overexpressing cells and tumors

Our team recently described the first reported small molecule inhibitors of IGF2BP2 (*92*). Taking advantage of our previous work, we tested four of these new compounds **(Figure 6A).** We generated dose‒response curves for these compounds in multiple PLK1-overexpressing breast cancer cell lines to determine their IC_50_. While compounds C4 and C9 inhibited the protein more effectively at lower concentrations **(Figure 6B)**, compound C4 showed the best *in vivo* pharmacokinetic (PK) profile of the tested IGFBP2 inhibitors. The highest exposures were observed in plasma, whole blood, blood cells, and urine after intravenous application in mice, during which the half-life was approximately 22 hours. In contrast, C6 and C9 had relatively low half-lives of only 1 hour, whereas C1 had a half-life of approximately 9 hours. Moreover, C4 had a high plasma C0 concentration of approximately 1.5 µg/ml after a relatively low dose of 1 mg/kg IV. Additionally, C4 and C1 exhibited low distributions (∼ 0.5 l/kg and 0.7 l/kg, respectively) and low clearances of ∼ 0.3 ml/min/kg and 0.9 ml/min/kg, respectively. Compounds C6 and C9 had moderate to low clearance rates of 25 and 10 ml/min/kg, respectively **(Figure 6C)**.

**Figure 6:**
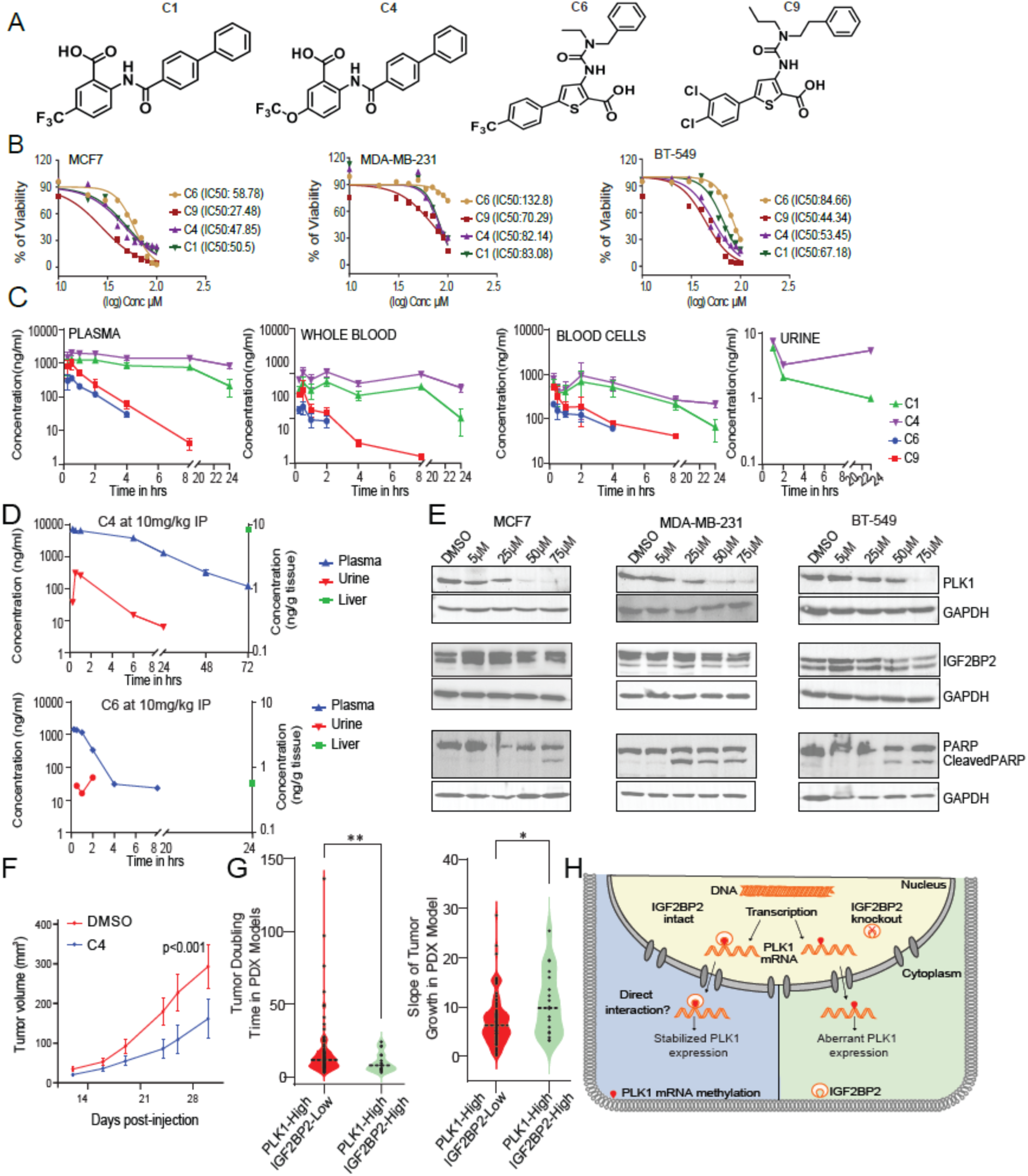
Pharmacological inhibition of IGF2BP2 reduces tumor growth. **A.** Chemical structures of the tested IGF2BP2 inhibitors. **B.** Dose‒response curves of the compounds (C1, C4, C6 and C9) that inhibited IGF2BP2 in PLK1-overexpressing cell lines. **C.** Concentrations of compounds C1, C4, C6 and C9 in plasma; whole blood; blood cells at 0.25, 0.5, 1, 2, 3, 8, and 24 h; and urine at 1, 2 and 24 hours after IV administration (1 mg/kg). **D.** Concentrations of compound C4 in plasma at 0.25, 0.5, 1, 6, 24, 48 and 72; in the liver at 72 and at 0.25, 0.5, 1, 6 and 24 hours after IP administration (10 mg/kg); concentration of compound C6 in plasma at 0.25, 0.5, 1, 2, 4, 8 and 24; in the liver at 24 and at 0.5, 1 and 2 hours after IP administration (10 mg/kg). **E.** Western blot showing the levels of PLK1, IGF2BP2 and PARP after treatment with an IGF2BP2 inhibitor (C4) in MCF7, MDA-MB-231 and BT-549 cells at doses ranging from 5 to 75 μM for 72 hours. GAPDH was used as a loading control. **F.** Effect of an IGF2BP2 inhibitor (C4) on tumor growth. MDA-MB-231 cells (2×10^6^) were injected into the mammary fat pads of immunodeficient female NOD-SCID mice. Treatment with the IGF2BP2 inhibitor at a dosage of 10 mg/kg was initiated the day after the injection of cells. The IGF2BP2 inhibitor and a matching volume of DMSO were dissolved in 2-hydroxypropyl-β-cyclodextrin and administered intraperitoneally (IP) for five consecutive days, followed by two days of rest, for a total of 30 days. Tumor volumes were measured with digital calipers in the control group treated with DMSO (n=10) and in the group treated with the IGF2BP2 inhibitor (C4) (n=10) every 3 to 4 days after starting treatment. The graph shows the mean tumor volume (± standard error of the mean). The group treated with the inhibitor showed a significant decrease in tumor volume compared to that of the control group (p<0.001, two-way ANOVA). **G.** PDX models were classified based on the expression levels of PLK1 and IGF2BP2. The tumor doubling times (left) and slopes of tumor growth (right) are presented for the two categories. **H.** Schematic model showing the regulation of PLK1 expression by IGF2BP2.

As C4 exhibited the best PK parameters out of the four compounds tested in the cassette PK study, we explored the intraperitoneal route at 10 mg/kg to enable administration over several days. Moreover, we tested C6 because it had a greater distribution volume of approximately 2 l/kg after IV administration, and we wanted to probe terminal compound levels in the liver as well **(Figure 6D)**. C4 showed sustained plasma levels above the IC_50_ (range of 47-82 nM in different cell lines), with a C_max_ of 7.1 µg/ml and a mean residence time of approximately 15 hours. No accumulation was observed as the plasma levels decreased until 72 hours. This finding suggested that a dosing interval > 24 hours could be used for efficacy studies of C4. Compound C4 was also found in the urine after up to 24 hours. Compound C6 had a C_max_ of approximately 1.5 µg/ml with a T_max_ of 0.4 h and a mean residence time of approximately 1.9 h. Compound C6 had terminal liver levels of approximately 32 ng/g tissue after 24 hours, whereas compound C4 still exhibited terminal liver levels of approximately 8.4 ng/g after 72 hours. Moreover, C4 had a bioavailability of approximately 32 %, whereas C6 had a bioavailability of approximately 38 % after IP administration. Based on these PK features, we chose C4 for further studies.

Using C4, we next assessed PLK1 and IGF2BP2 protein levels after 72 h of treatment with compound C4 at doses ranging from 5 μM to 75 μM. PLK1 protein levels consistently decreased after treatment with this inhibitor, although IGF2BP2 protein levels remained unaltered **(Figure 6E).** Finally, a PARP cleavage experiment was used to determine whether apoptotic pathways were activated in cells following treatment with C4. Indeed, we observed cleaved PARP in all tested cancer cells treated with higher doses of the compound **(Figure 6E).** To further evaluate the therapeutic potential of IGF2BP2 inhibition, we examined whether treatment with C4 would reduce tumor development in xenograft models. These experiments revealed that C4 administration suppressed tumor growth in triple-negative breast cancer xenografts generated with MDA-MB-231 cells **(Figure 6F).** Overall, these studies suggest that further optimization of IGF2BP2 inhibitors has high therapeutic potential, as the application of these compounds would allow us to effectively reduce PLK1 protein levels and ultimately inhibit PLK1-overexpressing cancer cells and tumors. To derive additional support for our findings, we took advantage of multiple previously described PDX models, representing different tissue types (*62, 63*). We evaluated the tumor growth patterns of 174 PDX models that were not subjected to any drug treatment and classified them into two groups based on the expression patterns of PLK1 and IGF2BP2. The first group represented patients with high PLK1 expression but low IGF2BP2 expression, while the second group represented patients with high PLK1 and IGF2BP2 expression. After analyzing the doubling time and the slope of tumor growth, we found that the PDX models in Group 1 had a significantly greater doubling time and significantly lower growth slope than did those in Group 2 (p<0.05, unpaired t test with Welch’s correction) **(Figure 6G)**. These results revealed that, in agreement with our observations in cultured cancer cells, IGF2BP2 also displays an SDL interaction and selectively suppresses the growth of PLK1-overexpressing tumors in PDX models. In summary, our work indicates that PLK1 levels can be modulated by IGF2BP2 and that its loss can decrease the growth of tumors with elevated PLK1 expression.

## Discussion

Multiple PLK1-targeting compounds have been identified and are currently being assessed in clinical trials (*93*). The most notable trials have been performed with BI2536, BI6727 (volasertib), and GSK461364A, which are all ATP-competitive inhibitors of PLK1. However, monotherapy with BI2536 was terminated due to a low objective response rate and poor survival (*94*), while GSK461364 was associated with a high incidence of venous thrombotic emboli in clinical studies (*95*). Volasterib, a derivative of BI2536, had initial success in gaining FDA breakthrough therapy status but has not shown significant promising results since then (*96*). New inhibitors of PLK1, such as TAK960 and NMS-P937, are still in their early stages (*93*). These findings indicate the importance of PLK1 targeting and highlight the challenges associated with its direct inhibition.

Here, we queried nearly the entire genome to identify genes that are selectively essential for the survival of PLK1-overexpressing cells. Our main goal is to develop an unbiased systematic approach to identify the optimal target for future therapeutic approaches. Therefore, we performed our initial genome-wide screen in an engineered model cell line. We used a PDX-derived cell line and a PDX xenograft model that naturally overexpresses PLK1 for further validation studies. Finally, we used direct capture Perturb-seq to query which identified SDL hits and eliminated most PLK1-overexpressing cancer cells. Thus, IGF2BP2 was identified by all the other approaches in our multipronged unbiased strategy. From a technical standpoint, we are also cognizant that genome editing tools have shortcomings associated with concerns about off-target effects (*97, 98*). Our combined use of orthogonal shRNAs and sgRNA reagents in validation approaches mitigated potential false positives associated with our screening and validation strategies. By using complimentary and independent validations with different gene-silencing and gene-editing techniques and model systems, the potential confounding effects were filtered out. We also demonstrated optimal strategies for applying pooled *in vivo* CRISPR screens by initially prioritizing our hits. Libraries can be reduced to a manageable size for both *in vitro* and *in vivo* experimental work while achieving meaningful results with a sufficient representation of guide RNAs.

Ten overlapping genes from the *in vivo* and *in vitro* PDX-based cohorts were identified and validated, several of which have a role in cancer progression and regulating stemness. For example, in addition to its role in centrosome clustering and cytokinesis, KIF5B has been shown to regulate the epithelial–mesenchymal transition (EMT) and stemness of cancer cells (*99*). Similarly, FDPS plays a role in maintaining tumor-initiating cells (TICs) in gliomas (*100*), and IGF2BP2 plays a role in maintaining TIC populations in both colon cancer and gliomas (*101, 102*). To further determine the best candidate gene in the context of PLK1 overexpression, we chose to use a single-cell CRISPR screen (Perturb-seq) by direct guide RNA capture. Since PLK1 overexpression can induce CIN, leading to intratumor heterogeneity, it is imperative to determine the efficiency of the SDL target at the single-cell level. This ultimately led to the shortlisting of four best candidates. Among these, IGF2BP2 is an N6-methyladenosine (m6A) reader (monitoring the presence of a methyl group at the *N*6 position of adenines in poly(A)^+^ RNA) that has been shown to bind m6A-methylated mRNA molecules to enhance their stability and translation (*103*). As the mRNA of PLK1 has been previously reported to be m6A methylated (*104*), we speculate that one of the potential mechanisms by which IGF2BP2 can regulate PLK1 expression could be via the stabilization of the m6A-methylated mRNA of PLK1 **(Figure 6H)**. Consistent with this notion, while our work was in progress, a recent study showed that IGF2BP2 binds to m6A in the 3’ untranslated region of PLK1 and is involved in stabilizing PLK1 expression (*105*). Interestingly, the loss of IGF2BP2 not only affects PLK1 mRNA but also reduces the abundance of the PLK1 protein in multiple models, including MDA-MB-231 breast cancer cells, where its inactivation does not decrease the abundance of PLK1 mRNA. Taken together, these data suggest that IGF2BP2 operates in cancer cells in a context-dependent manner but that its SDL relation with PLK1 relies on PLK1 suppression. This PLK1 inactivation eventually causes the selective elimination of PLK1-overexpressing cancer cells, which depend on elevated PLK1 activity.

Interestingly, both PLK1 and Aurora kinases are overexpressed within the same single cells, and loss of IGF2BP2 appears to eliminate most of these single cells. Aurora kinases are a family of therapeutic targets that are highly overexpressed in cancer cells and functionally linked to PLK1 (*24*). Although there are more than 20 Aurora kinase inhibitors in different stages of clinical development, prolonged treatment has been shown to lead to the development of drug resistance (*106*). Targeting IGF2BP2 could potentially eliminate cells that co-express both PLK1 and Aurora kinases. Although the concentration of C4 used to inhibit IGF2BP2 is relatively high, it is also important to note that at 10mg/kg IP, C4 reached sufficient concentrations in plasma to warrant further efficacy studies. Therefore, we expect C4 to have the potential to reach efficacy in the nanomolar range in combination therapies with good pharmacokinetic properties.

While we have focused on IGF2BP2, further exploration of the other three candidates is needed to determine their SDL relationships with PLK1. For example, CRB3 and TJP3 are involved in the establishment of tight junctions (*107, 108*) and may impact cytokinesis, as PLK1 also plays a key role in this process (*109*). In fact, among the pleiotropic defects caused by PLK1 overexpression (*20, 23–26*), cytokinesis failure and abscission are the most common defects, resulting in aneuploidy (*22*). Further studies to explore the role of these proteins might also point toward new therapeutic avenues. On the other hand, DPP9 is a peptidase that degrades most cytosolic proline-containing peptides (*110*) and has been shown to interact specifically with SUMO1 (*111*). The function of DPP9 may facilitate mitotic entry by affecting ubiquitination of the transcription factor Forkhead box protein M1b (FoxM1b), a known substrate of PLK1 (*112*). Thus, our work has led to several testable models for discovery science, and we expect the research community to benefit from our extensive validation strategies.

Our systematic integration of multiple unbiased platforms identified a large resource of PLK1-SDL hits that can be targeted to suppress the proliferation of PLK1-overexpressing cells. As PLK1 overexpression can lead to cellular heterogeneity, it is also necessary to confirm whether suppression of its SDL partner(s) truly eliminates PLK1-overexpressing individual cells. Direct-capture Perturb-seq revealed IGF2BP2, the loss of which affects PLK1 protein levels through genetic or pharmacological inhibition. In this regard, we are actively working on optimizing our novel IGF2BP2 inhibitors to use them effectively in preclinical studies, with the ultimate goal of progressing them into clinical trials.

## Acknowledgments

We thank members of the Vizeacoumar and Freywald laboratories for their insights and comments on the manuscript. We thank Dr. Alana L. Welm, Co-Director, Cell Response and Regulation Program, Huntsman Cancer Institute, University of Utah, for providing the HCI-010 PDX and for the related technical advice. We thank Dr. Laurent Creancier, Pierre-Fabre, France, for providing HCT116 inducible PLK1 cells. We also thank Janine Schreiber and Jennifer Wolf for excellent technical assistance. The studies were approved by the Animal Ethics Research Board of the University of Saskatchewan and appropriate regulatory authorities (# 20150067) and the ethical board of the Niedersächsisches Landesamt für Verbraucherschutz und Lebensmittelsicherheit, Oldenburg, Germany. **Financial support:** Canadian Institutes of Health Research operating grants PJT-156309 and PJT-156401 (FJV, AF). Cancer Research Society operating funds 2017-OG-22493 (FJV, AF). Be Like Bruce Foundation (FJV, AF). Deutsche Forschungsgemeinschaft #453246190 (AKK, ME). Canada Foundation for Innovation CFI-33364 (FJV). Cancer Foundation of Saskatchewan (FJV). Lisa Rendall Fellowship (CEC). College of Medicine, University of Saskatchewan (FSV). Mitacs Globalink Fellowship (LK). SHRF Fellowship (RD). University of Saskatchewan CoMGRAD Award (VM, HP, HE). Canadian Institutes of Health Research - Canada Graduate Scholarships – Master’s (JDWP). University of Saskatchewan Health Science Graduate Scholarship (JDWP). Cancer Research Society – Doctoral Research Award (JDWP). Canadian Institutes of Health Research - Canada Graduate Scholarships – Doctoral FBD-187665 (VM).

## Author contributions

Conceptualization: F.S.V., M.E., A.K.K., A.F., F.J.V; Investigation: C.E.C., F.S.V., Y.Z., L.K., P.G., V.M., H.D., J.P., A.G., R.G., C.D., S.B., K.W., Y.W.1, F.K., S.M., A.C., T.K., B.G., E.Z., K.R., P.W., L.G., H.P., A.M.M., K.K.B., H.E., R.D., O.A., A.D., T.F., E.P.M., J.S.L., B.T.; Writing, review & editing: All authors; Funding Acquisition: A.F., A.K.K., M.E., F.J.V.

## STAR Methods

### Resources availability

#### Lead contact

Further information and requests for reagents and resources should be directed to and will be fulfilled by the lead contact, Franco Vizeacoumar.

#### Materials availability

This study did not generate new unique reagents.

#### Data and code availability

The data files were uploaded to the Gene Expression Omnibus (GEO) database, and the SuperSeries accession number is GSE203242. This reference consists of two SubSeries for both the raw microarray data (.CEL files) for the shRNA screen GSE203237 and the deep sequencing data from the CRISPR screen (.fastq.gz files) GSE203240. All other data reported in this paper will be shared by the lead contact upon request. This paper does not report original code. Any additional information required to reanalyze the data reported in this paper is available from the lead contact upon request.

### Experimental model and study participant details

#### Cell lines and cell culture

All cell lines used in this study are listed in Supplementary Table S11. Cell lines were incubated at 37°C in 5% CO2. All cell lines were purchased from Cedarlane labs, unless otherwise indicated (Burlington, Ontario, Canada), a Canadian distributor for American Type Culture Collection (ATCC) or from Millipore Sigma. Cell lines purchased from ATCC/Sigma were passaged for less than three months at a time following resuscitation; therefore, no additional authentication was performed. Mycoplasma tests were routinely conducted.

#### Mouse models, tumor xenograft and pharmacokinetic studies

Mice were housed 5 per cage at 23-25°C in a humidity-controlled colony room, maintained on a 12h light/dark cycle (08:00 to 20:00 light on), with standard food and water provided *ad libitum* and environmental enrichments. All animals were handled in accordance with the approved protocols by the University of Saskatchewan Animal Research Ethics Board (AREB). All mice used in this study were NSG (NOD.Cg-Prkdscidll2rg) and littermates within the same cage were randomly allocated to experimental or control groups.

The mice used in the present study were from our established colony of immunodefficient female NOD mice. Cg-*Prkdc^scid^Il2rg* (NOD-SCID) mice were obtained from the Laboratory Animal Services Unit (LASU), University of Saskatchewan. The cells were trypsinized and resuspended in ice-cold PBS. MDA-MB-231 cells were injected into the inguinal mammary fat pads of 6- to 10-week-old female NOD-SCID mice (2×10^6^ cells per mouse). DMSO (5%) and C4 (10 mg/kg) were dissolved in PBS containing 5% 2-hydroxypropyl-β-cyclodextrin (HP-β-CyD) as an excipient and injected intraperitoneally five times per week for four weeks (DMSO, n = 10; C4, n = 8) in a total volume of 100 µL.

Treatment with an IGF2BP2 inhibitor (Compound 4) was initiated the day after the injection of MDA-MB-231 cells. Injections were given for five consecutive days, followed by two days of rest, for a total of 30 days. Tumors were measured every three to four days using a digital caliper, and the tumor volume was calculated using the formula (A*B^2^)/2, where A and B represent the long and short diameters of the tumor, respectively.

For pharmacokinetic studies, 4-week-old outbred male CD-1 mice (Charles River, Germany) were used. The animal studies were conducted in accordance with the recommendations of the European Community. All animal procedures were performed in strict accordance with the German regulations of the Society for Laboratory Animal Science (GV-SOLAS) and the European Health Law of the Federation of Laboratory Animal Science Associations (FELASA). Animals were excluded from further analysis if sacrifice was necessary according to the human endpoints established by the ethical board. All experiments were approved by the ethical board of the Niedersächsisches Landesamt für Verbraucherschutz und Lebensmittelsicherheit, Oldenburg, Germany. Compounds C1, C4, C6 and C9 were dissolved in 1.6 % DMSO, 20 % PEG400, 20 % Tris (1 % pH 9.0) and 58.4 % 0.9 % isotonic NaCl solution. Mice were administered C1, C4, C6 and C9 in a cassette PK format at 1 mg/kg IV per compound. Approximately 20 μl of whole blood was collected serially from the lateral tail vein at 0.25, 0.5, 1, 2, 4, and 8 h post administration. After 24 h, the mice were euthanized, and blood was collected from the heart. Whole blood was collected into Eppendorf tubes coated with 0.5 M EDTA. One part was immediately spun down at 13.000 rpm for 10 min at 4°C. The plasma was transferred into a new Eppendorf tube and then stored at −80°C until analysis. Additionally, blood cells and whole blood were preserved and used for analysis. Furthermore, C4 and C6 were tested in a single-dose PK study at 10 mg/kg IP. C4 and C6 were dissolved in 10 % DMSO and 90 % Tris (1 % pH 9.0). Approximately 20 μl of whole blood was collected serially from the lateral tail vein at 0.25, 0.5, 1, 6, 24 and 48 h post administration for C4 and at 0.25, 0.5, 1, 2, 4 and 8 h post administration for C6. After 72 h for C4 mice and after 24 h for C6 mice, blood was collected from the heart, and whole blood was collected into Eppendorf tubes coated with 0.5 M EDTA and immediately spun down at 13,000 rpm for 10 min at 4°C. The plasma was transferred into a new Eppendorf tube and then stored at −80°C until analysis. Moreover, spontaneous urine was collected from the cassette and from the single-dose studies. Liver samples were homogenized in isotonic sodium chloride solution using a Polytron® homogenizer.

All PK plasma samples were analyzed via HPLC‒MS/MS using an Agilent 1290 Infinity II HPLC system coupled to an AB Sciex QTrap6500plus mass spectrometer. First, a calibration curve was prepared by spiking different concentrations of C1, C4, C6 and C9 into mouse plasma, mouse whole blood, mouse blood cells and mouse urine from CD-1 mice. Caffeine was used as an internal standard. In addition, quality control samples (QCs) were prepared for C1, C4, C6 and C9 in the respective matrices. The same extraction procedure was used: 7.5 µl of a plasma or whole-blood sample, 5 µl of a blood cell sample + 7.5 µl of isotonic sodium chloride solution or 10 µl of a urine sample (from the calibration samples, QCs or PK samples) was extracted with 37.5 µl of methanol containing 12.5 ng/ml caffeine as an internal standard for 10 min at 2000 rpm on an Eppendorf MixMate® vortex mixer. Then, the samples were spun down at 13,000 rpm for 10 min at 4 °C. The supernatants were transferred to standard HPLC glass vials. 50 µl of a homogenized liver sample (adjusted to a final concentration of 300 mg/ml; calibration samples, QCs or PK samples) was extracted with 50 µl of methanol containing 12.5 ng/ml caffeine as an internal standard for 10 min at 800 rpm on an Eppendorf MixMate® vortex mixer. Then, the samples were spun down at 4,000 rpm for 10 min at 4°C. The supernatants were transferred to 96-well V-bottom Greiner plates and sealed. The HPLC conditions used were as follows: column: Agilent Zorbax Eclipse Plus C18, 50×2.1 mm, 1.8 µm; temperature: 30°C; injection volume: 1 µl; flow rate: 700 µl/min; solvent A: water + 0.1 % formic acid; solvent B: acetonitrile + 0.1 % formic acid; gradient: 99% A at 0 min and until 0.1 min 99 % - 0% A from 0.1 min to 4.0 min, 0 % A until 4.5 min, 0 % - 99 % A from 4.5 min to 4.7 min. The mass spectrometric conditions were as follows: scan type: MRM, positive and negative mode; Q1 and Q3 masses for caffeine, C1, C4, C6 and C9 can be found in **Table S10**; and peak areas of each sample and the corresponding internal standard were analyzed using MultiQuant 3.0 software (AB Sciex). PK parameters were determined using a noncompartmental analysis with PKSolver (*61*).

### Methods details

#### Transfections and transductions

Lentiviral particles were generated by transfecting HEK293T cells with psPAX2, pMD2.G, and a pLKO.1-shRNA 9:1:10 plasmid mixture was generated using Xtremegene 9 transfection reagent (Roche) and Opti-MEM media (Gibco). The media was replaced with DMEM containing 2% (w/v) bovine serum albumin 18 hours post-transfection, and lentivirus-containing media was harvested at 48- and 72-hours post-transfection. Transfections with siRNA for Hs578Bst were carried out using RNAiMax transfection reagent (Life Technologies) and Opti-MEM to a final siRNA concentration of 50 nM. Cas9-expressing HCI-010 cells were generated by transducing HCI-010 cells with Cas9-blast lentiviral particles (Addgene #52962), replacing the lentiviral media after 24 hours, and adding 5 µg/mL blasticidin 48 hours after transduction for 21 days. The antibodies GAPDH (sc-25778) and PLK1 (sc-5585) were obtained from Santa Cruz, Cas9 (ab191468), IGF2BP2 (ab128175) from Abcam, tubulin (#T902) from Merck, and PARP (9542S) from Cell Signaling.

#### Genome-wide pooled shRNA screening and data analysis

Screening and microarray scoring were performed as previously described (*50*). HCT116-PLK1 cells were transduced at an MOI of 0.3, and 24 hours after transduction, the cells were treated with 2 µg/mL puromycin for 48 hours. Puromycin-selected cells were divided into two populations, and one population was induced with doxycycline (PLK1-IN) on day 0. The cells were passaged over 16 days with 200x hairpin representation, and samples were collected at days 0, 8, and 16. Genomic DNA was extracted at each timepoint, and shRNA sequences were PCR-amplified at 98°C for 3 minutes, followed by 98°C for 10 seconds, 55°C for 15 seconds, and 72°C for 15 seconds for 29 cycles. Amplified hairpins were digested with XhoI, and the stable half hairpin gel was extracted, purified, and subjected to probe hybridization on UT-GMAP 1.0 microarrays. The signal intensities for the microarray were normalized via quantile normalization, and shRNAs with signals below the background (i.e., a log_2_ scale of less than 8) at the initial timepoint T0 and signals below 0 at timepoints T8 and T16 were removed prior to further analyses to compute the fitness score. The weighted differential cumulative change (WDC_h_) for each shRNA between the doxycycline-induced cells (PLK1-IN) and its isogenic uninduced cells (PLK1-UN) was calculated for each consecutive timepoint using the following formula:

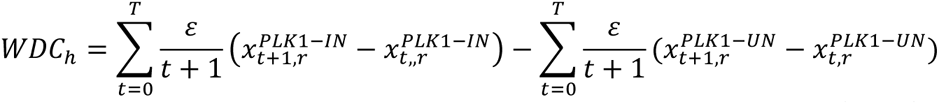

where 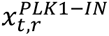 is the normalized signal intensity of the PLK1-IN cells at time point *t* ∈ (0, . . *T*) in replicates *r* ∈ (1. . *N*).

Similarly, 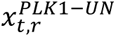 represents PLK1-UN cells. ε is a constant that determines the weight between each time point so that shRNA drops at earlier time points are ranked before the shRNA drops at later time points. The WDC_gene_ gene was computed as the average of the two shRNAs with the most negative values for that gene using the formula below.

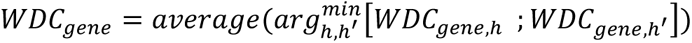

To identify shRNAs and their corresponding genes that are significantly different between PLK-IN and PLK1-UN cells, Student’s t test was used in combination with the permutation test p value by estimating the frequency of randomized, shuffled WDCs with more negative values in comparison with the observed gene-level WDC value, as previously described (*50*). Bayesian analysis of the gene essentiality algorithm was used to evaluate the performance of the screens (*51*).

#### Computational analysis and datasets

Reactome pathway enrichment was performed by using the Reactome database to assign genes to pathways (https://reactome.org/) (*52*), after which gene enrichment analysis was performed on the query gene set of 960 PLK1 screening hits using the idep software package (*53*). Gene expression analysis was performed using The Cancer Genome Atlas database downloaded from the Genomic Data Commons data portal (https://portal.gdc.cancer.gov/) for 33 different cancer types. The RNA-Seq Expectation Maximization (RSEM)-normalized mRNA expression data were log2 transformed, and the Spearman-rank correlation between PLK1 and each of the 960 SDL hits was calculated for each of the 33 different malignant patient samples. The resulting correlation coefficient (r) was subsequently clustered by calculating the Euclidean distance between different genes.

The gene expression profile from the Cancer Cell Line Encyclopedia database (*54*) was used to rank cell lines according to PLK1 expression to classify them into two groups of cell lines that over- or under-express PLK1. For the essentiality datasets from Project Achilles (*55*) and Marcotte *et al*.(*56*), the top 25% and bottom 25% of the ranked cell lines were assigned as PLK1-overexpressing and under-expressing respectively. To determine if a PLK1 screening hit was classified as essential or non-essential, the fold change in the essentiality scores between the PLK1-overexpressing and PLK1-underexpressing groups, as well as the p value, was calculated using the Mann‒Whitney U test.

ToppGene candidate gene prioritization (*57*) was performed using “PLK1” as the training gene set and the list of 960 PLK1 screening hits as the test gene set. The ToppGene database was used to compile functional annotations and network analysis results from Gene Ontology (GO) for molecular function, biological process, cellular component, gene expression, pathway, protein domain, transcription factor-binding site, miRNA target, drug–disease interaction, disease– drug interaction, and interaction information published in the NCBI search engine PubMed. Gene expression data from the TCGA were used to divide patients into two groups based on PLK1 expression and PLK1 screening hit expression for each of the 960 PLK1 screening hits. The two groups are illustrated in **Figure 2D**. The median gene expression values for PLK1 and for the screening hits were determined and used as the cutoff values to divide samples into PLK1 low and PLK1 high. TCGA clinical data were subsequently subjected to standard Kaplan‒Meier analysis between patients who exhibited natural SDL and those who did not.

Drug response analysis was performed by using the Genomics of Drug Sensitivity in Cancer (GDSC) dataset. The GDSCs are involved in gene expression and drug response to various drugs in several cancer cell lines. We grouped the cell lines based on the expression of PLK1 into the top 1% and bottom 1%. Then, we computed the p value for the difference in the p value between the PLK1-high and PLK1-low cells using the Mann‒Whitney U test for the IC50s of the inhibitors of the SDL hit present in the GDSC.

#### Pooled in vivo CRISPR screening

Mice were housed under sterile conditions at the University of Saskatchewan, and all animal protocols were reviewed and approved by the University of Saskatchewan Animal Research Ethics Board. Digital caliper measurements were used to monitor tumor size, and tumor volume was calculated by the formula A/2*B2 (where A and B are the long and short diameters, respectively). The sgRNA sequences used in the imaging experiment were pooled and used to generate a pooled lentiviral library. HCI-010 and HCI-010+Cas9 PDX-derived cells (*58*) were transduced at an MOI of 0.3, and after 24 hours, they were selected using 2 µg/mL puromycin. After 48 hours of selection, 3 million viable cells mixed 1:1 with Matrigel (Corning) in a total volume of 100 µL were injected into the mammary fat pads of 6- to 8-week-old female mice and allowed to grow for 3 weeks. The tumors were harvested, and flash frozen in liquid nitrogen. Genomic DNA was extracted using a mortar and pestle and a DNA Blood Maxi Kit (Qiagen). A sequencing library was constructed as described previously (*59*) with slight modifications to the primers of the first PCR to match the vector backbone (*60*) (forward primer 5-caaaatacgtgac gtagaaagtaataatttcttgggtag-3’ and reverse primer 5’-gcgtaaaattgacgcatgtgttttatcggtctgtatatcgag-3’).The fastq files were aligned to the sequence library using the Bowtie2 alignment package. The alignment with scores above the default threshold score was mapped to the library. The count matrix data were analyzed as described in the pooled screen analyses above.

#### Microscopy and imaging

HCI-010 and HCI-010+Cas9 cells were seeded in 96-well low-attachment plates at a concentration of 10 000 cells per well with 8 µg/mL polybrene and 100 µL of sgRNA-lentiviral targeted pool (2 sgRNAs for each target gene, one gene per well) in a 200 µL total volume. After 24 hours, the media was replaced with media containing 2 µg/mL. After 48 hours of puromycin selection, 4 sites per well were imaged with a 4x objective using the brightfield setting for several days (Day 0, Day 4 and Day 6). After imaging, the images were analyzed using the MetaXpress custom module, where the cells were carefully identified by eliminating any artifacts. The custom algorithm was also used to measure the area, perimeter and number of cells per well. The increase in cell confluency each day was compared to baseline (Day 2), and HCI-010+Cas9 was subsequently compared to HCI-010 to determine whether the gene knockouts caused a decrease in cell proliferation.

#### Direct capture of guide RNA using a Perturb-seq screen

The 10x Genomics Chromium Next GEM Single-cell 3’ Reagent Kit-compatible sgRNA library, which included 130 sgRNAs (targeting 65 genes), 15 nontargeting controls, and 16 positive controls, was generated by MilliporeSigma. Samples for direct-capture single-cell Perturb-seq were prepared by transducing MCF7-Cas9 cells with single-cell perturb-seq sgRNA library lentiviral particles and harvested 2 days after 48 hrs of puromycin selection. The 3’Gene Expression Library and CRISPR Screening Library were constructed following the instructions of the Chromium Next GEM Single-cell 3’Reagent Kit v3.1 (10x Genomics). Both libraries were sequenced through the NovaSeq S1 Flow Cell platform (Illumina). The raw data from the NovaSeq platform were extracted as fastq.gz files. These files were mapped to the GRC38 reference genome and aligned using the cell ranger (v6.1.2) count algorithm run on 32 core cluster computing datasets. The samples were split into three batches, with the cell expectancy estimated to be approximately 10,000 cells per batch. The three batches were pooled together by using the cell ranger aggregate algorithm. Upon further investigation, it was found that several cells had more than one sgRNA. Hence, cells with a single sgRNA were curated and recomputed using the cell Ranger reanalysis pipeline. The resulting output was then imported into Loupe Browser (v.6.0) to generate various visualization outputs.

#### Colony formation

The cells (1×10^3^/well for MCF7 cells, MDA-MB-231 cells, 3×10^3^/well for HCC1143 cells, 5×10^3^/well for BT-549 cells) were seeded in 6-well plates and incubated at 37°C under 5% CO_2_ for 10 days, during which the media was replaced every 3 days. The plates were then washed with PBS, fixed with 4% formaldehyde and stained with 0.5% crystal violet. Colonies were counted using ImageJ software, and images of the colonies were scanned.

#### Genomic cleavage detection

Primers for genomic cleavage detection were designed for the region around the sgRNA target sequence with the forward primer ∼200 bp upstream and the reverse primer ∼300 bp downstream using Primer3 v. 0.4.0 (http://bioinfo.ut.ee/primer3-0.4.0/). The primer sequences are given in **Supp. Table S9.** The cells were lysed, the sgRNA target region was PCR amplified, and mismatched enzymes were digested with T7 endonuclease using a GeneArt Genomic Cleavage Detection Kit according to the manufacturer’s directions (Invitrogen).

#### Droplet digital PCR

cDNA templates for droplet digital PCR were prepared with a High-Capacity cDNA Reverse Transcription Kit (Applied Biosystems). The 20 µL quantitative polymerase chain reaction was performed using 10 µL ddPCR™ Supermix for Probes (Bio-Rad), 1 µL of target gene probe (*IGF2BP2*: FAM-MGB, Hs00538954_g1, *PLK1*: FAM-MGB, Hs00983227_m1, Thermo Scientific), 1 µL of reference probe (*GAPDH*: VIC-MGB, Hs02786624_g1, Thermo Scientific), and 50 ng of cDNA template. Droplets were generated from a QX100 ddPCR Droplet generator (Bio-Rad Laboratories). Quantitative PCR was performed with a CFX96 Real-Time PCR Thermocycler (Bio-Rad Laboratories) using the following program: 95℃ 10 minutes, 45 cycles of: 94℃ 30 seconds, 60℃ 90 seconds, 98℃ 10 minutes after 45 cycles, 4℃ on hold. The signals were detected with a QX100 ddPCR Droplet Reader (Bio-Rad Laboratories), and absolute quantification was performed with QuantaSoft software (Bio-Rad Laboratories).

#### Cell viability assay

The viability-inhibiting activity of the chemical compounds was assessed using a resazurin reduction assay. The MDA-MB-231, MCF7 and BT-549 cell lines were treated with increasing concentrations of the drugs (from 5 to 75 µM) and incubated for 72 h before resazurin (R&D Systems) was added. The fluorescence was detected by a Varioskan LUX Plate Reader (Thermo Fisher). Each experiment was conducted thrice, each with six replicates. Dose–response curves were generated, and the 50% inhibitory concentrations (IC_50_) were calculated using GraphPad Prism 9.

#### Proliferation assay in tumorspheres

The cells were seeded into 24-well Ultra-Low attachment plates (4×10^3^/well) in complete MammoCult medium (StemCell) and allowed to propagate in tumourspheres for 7 days. For each replicate of the IGF2BP2-KD cell line and corresponding control (shRFP), tumourspheres from 4 independent wells were combined and dissociated using mechanical dissociation (10 replicates of each sample were pipetted on P1000), and proliferation was assessed by cell counting using a hemocytometer. Each sample was assessed in triplicate, and three independent experiments were performed. Pictures were taken using an EVOS fluorescence Microscope (Invitrogen). Images were converted to 8-bit binary images and analyzed using ImageJ software.

#### Analyses of PDX models

The tumor growth patterns of 174 PDX models from various tissue types, such as breast, colorectal, non-small cell lung carcinoma, melanoma and pancreas tissue types, that had matching RNA-Seq data were taken from previously published studies (*62*). The samples of these 174 PDX models that were not exposed to any drug treatment were chosen for further analysis. The slope of the tumor growth curve was calculated using the Xeva package (v.1.99.20) (*63*). The doubling time of the tumor was taken from the original publication (*62*). The grouping of the PDX models was based on their RNA-Seq expression in fragments per kilobase of transcript per million mapped fragments (FPKM) units. A FPKM value <20 was considered to indicate low expression, and a value >30 was considered to indicate high expression of both PLK1 and IGF2BP2, as most PDX samples had high expression of PLK1. Accordingly, the PDX models were grouped into two categories: PLK1 high expression and IGF2BP2 low expression (representing SDL) and PLK1 and IGF2BP2 high expression. Finally, the slope and doubling time of each group were plotted in violin plots, and their significance was calculated using an unpaired t test with Welch’s correction.

#### Statistical Analysis

GraphPad prism software was used for all tests. A one-tailed unpaired Student’s t-test was used for analysis. Data are presented as mean ± standard deviation (SD). Significance of our results was determined by setting p < 0.05 and error was reported as plus or minus the standard deviation (SD). The non-parametric Mann–Whitney U test was used to compare two groups. Survival analyses were performed using the Kaplan–Meier estimator with the non-parametric log-rank test to measure the equity of strata. For xenograft work, unpaired heteroscedastic t-test was used for statistical significance.

## Supplemental information

**Fig. S1.**
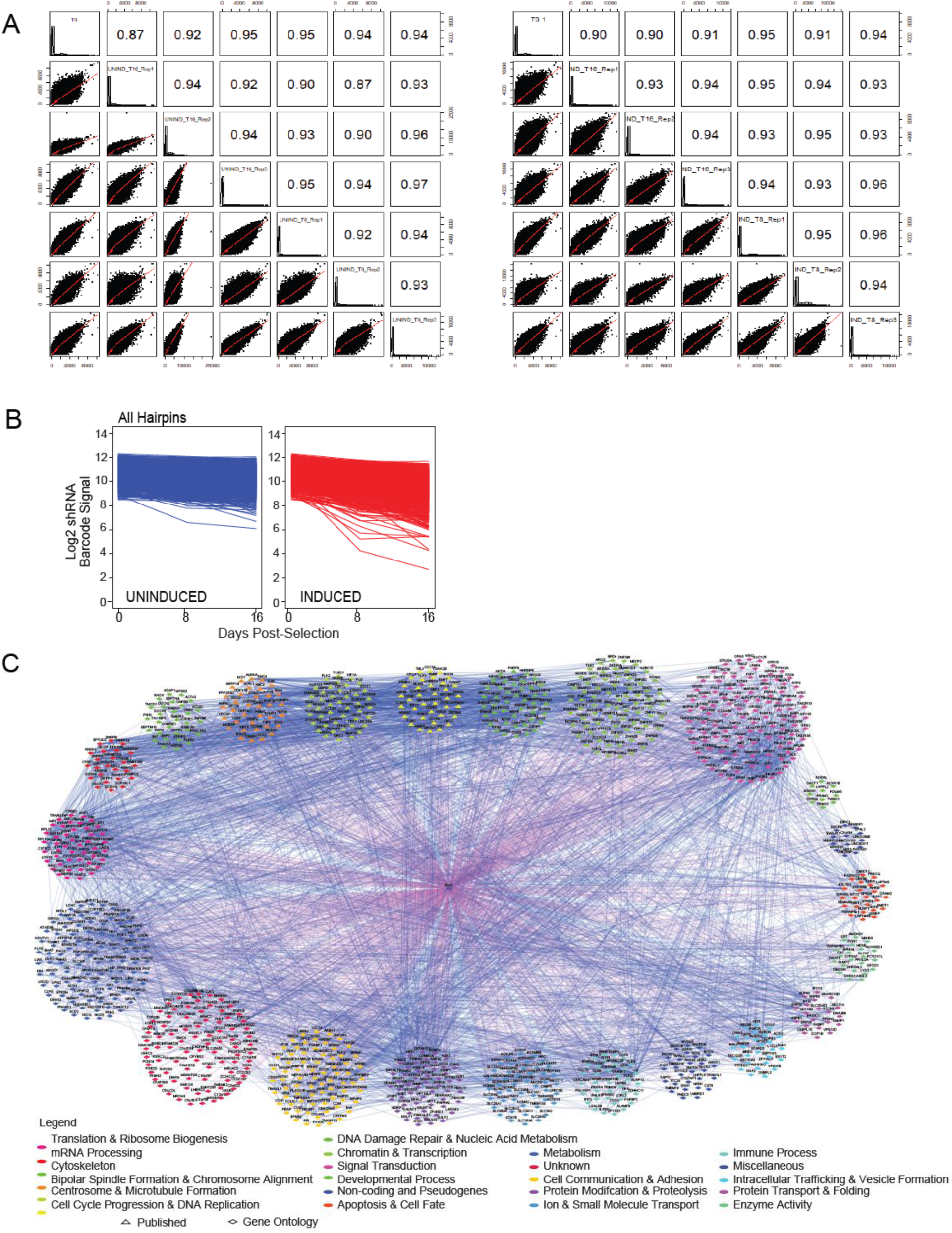
**A.** Correlations between replicates of uninduced and induced samples according to the genome-wide shRNA screens. Correlations between multiple timepoints are presented. The left panel shows the correlation between uninduced samples, and the right panel shows the correlation between induced samples. **B.** Magnitudes of dropouts between uninduced and induced samples at different time points for each hairpin from the genome-wide shRNA screen are plotted. **C.** Cytoscape network of all 960 SDL hits identified by genome-wide shRNA screens. Each node represents a hit from the screen. The genes are color coded based on their gene ontology. The blue edges represent previously published interactions downloaded from the STRING database showing crosstalk among the SDL hits.

**Fig. S2.**
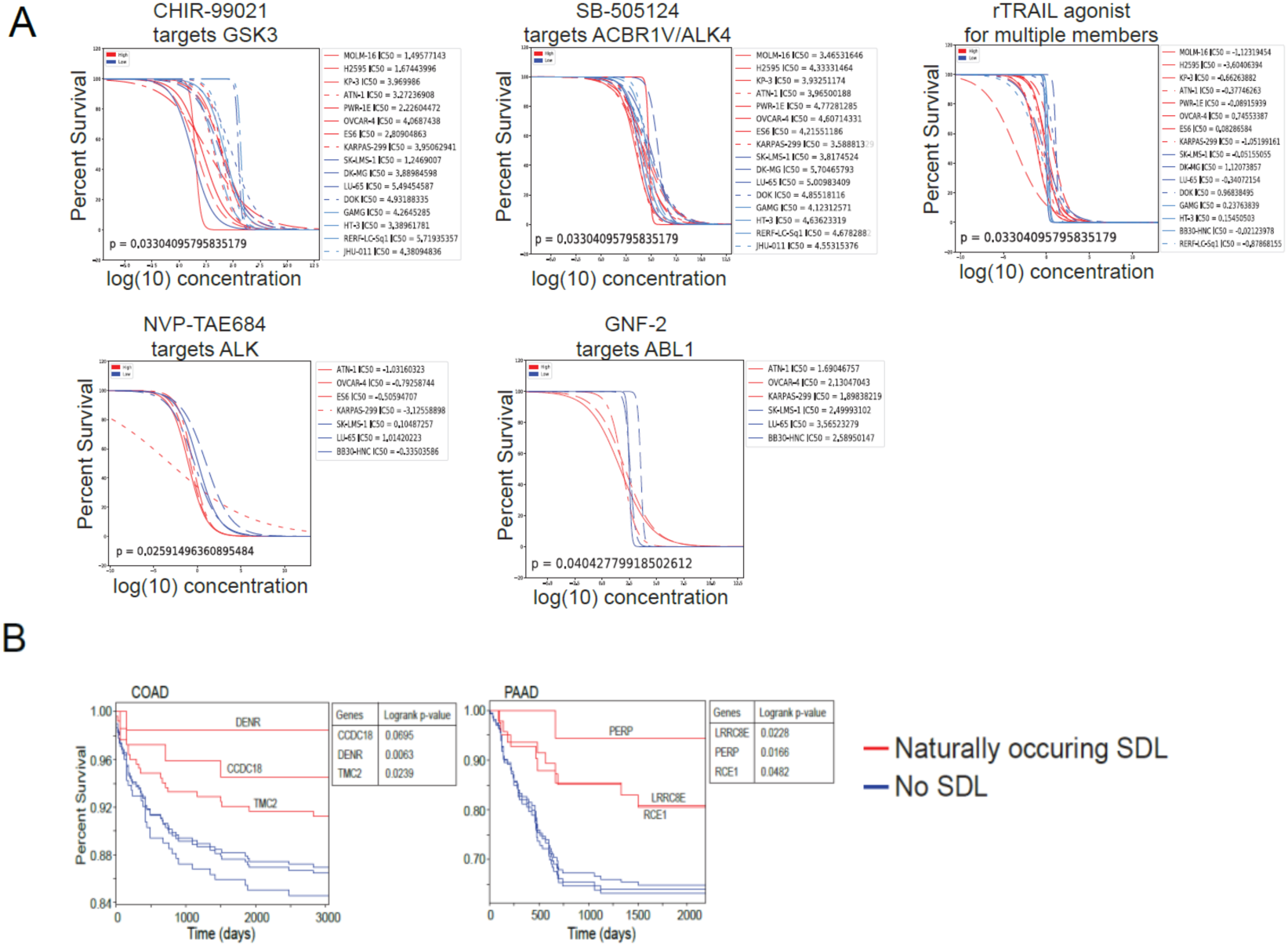
**A.** IC50 curves of drug inhibitors of some of the SDL hits identified in the screen. The red sigmoidal curve represents the IC50 curve for PLK1-overexpressing cell lines, and the blue sigmoidal curves represent the IC50 curves for cell lines with low PLK1 expression. **B.** Few representative examples of Kaplan‒Meier plots for colon and pancreatic cancer patients displaying natural SDL expression patterns are presented.

**Fig. S3.**
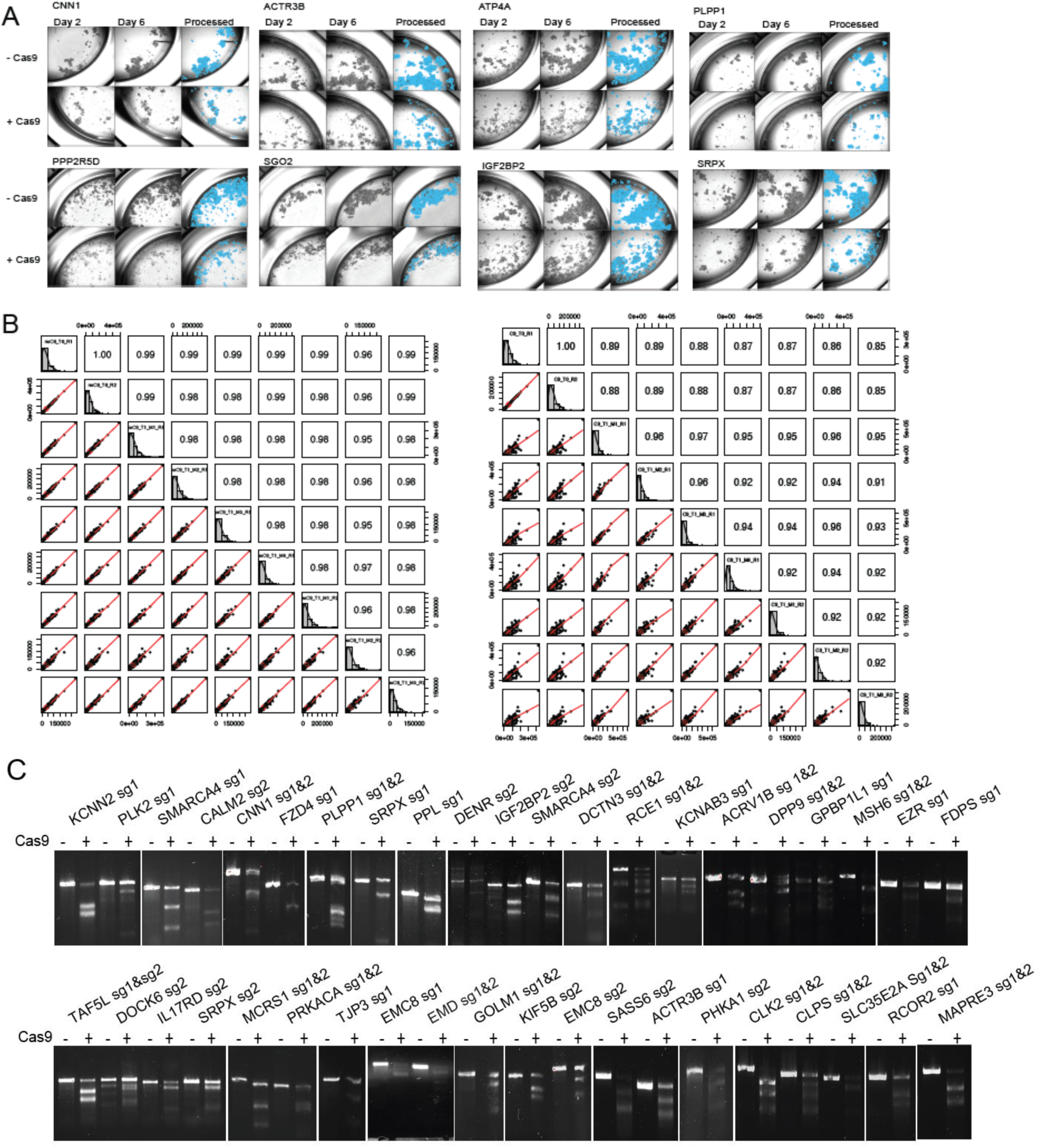
**A.** Representative images acquired using automated imaging over time for different PLK1-SDL candidate knockouts in Cas9^-^ and Cas9^+^ HCI-010 cells. The MetaXpress object masking overlay is shown in blue for the day 6 images. **B.** Correlation plots between replicates of the *in vivo* pooled CRISPR screen. The left panel shows the Cas9-negative samples, and the right panel shows the Cas9-positive samples. **C.** Representative cleavage assay confirming the individual knockouts.

**Fig. S4.**
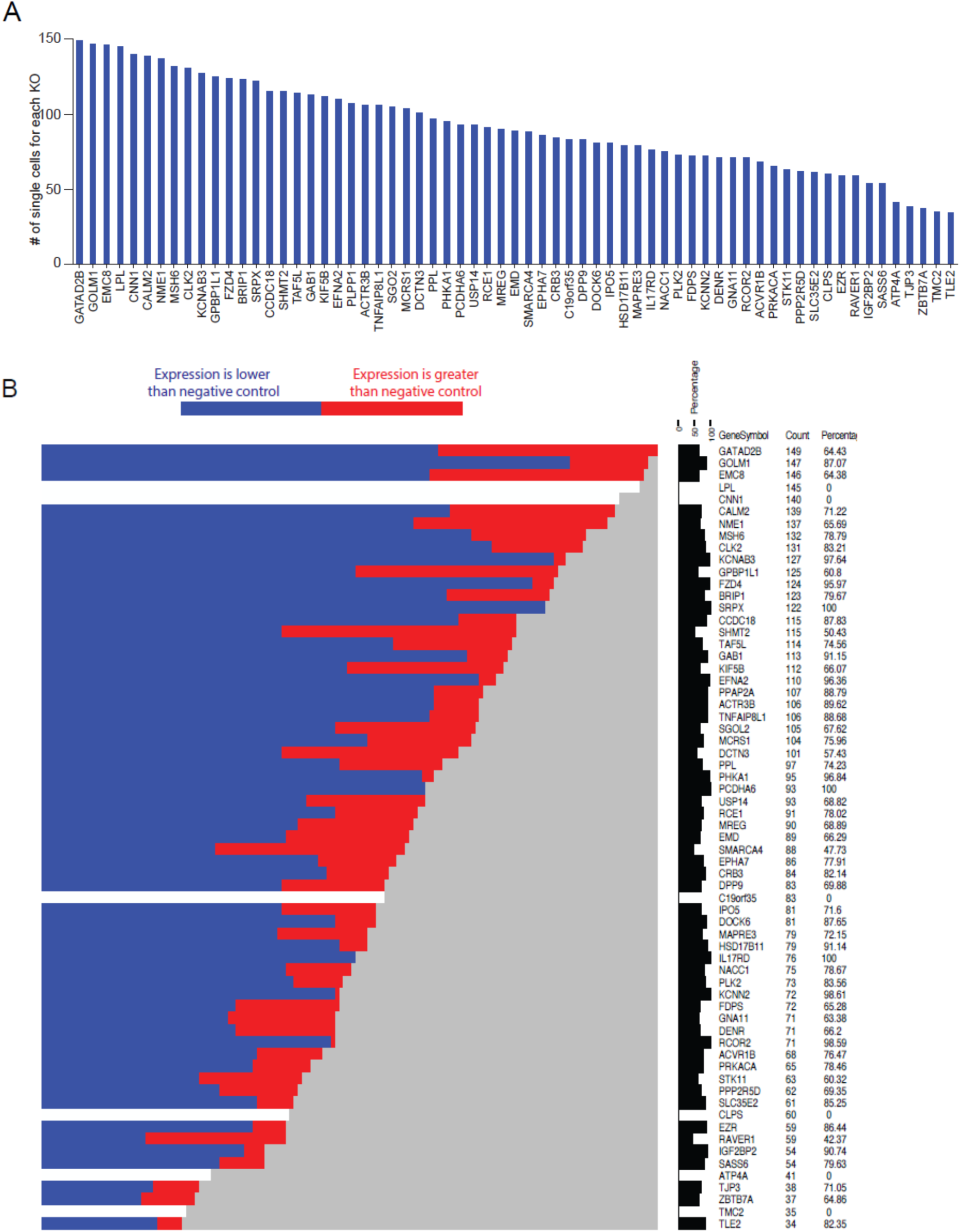
**A.** Bar graph showing the number of cells with each individual knockout analyzed via single-cell direct capture Perturb-seq. **B.** Knockout efficiency in the single-cell CRISPR screen was confirmed by comparing the expression of the corresponding target gene between cells from negative controls and knockouts. The percentage of knockout efficiency for each target is presented in addition to the heatmap.

**Fig. S5.**
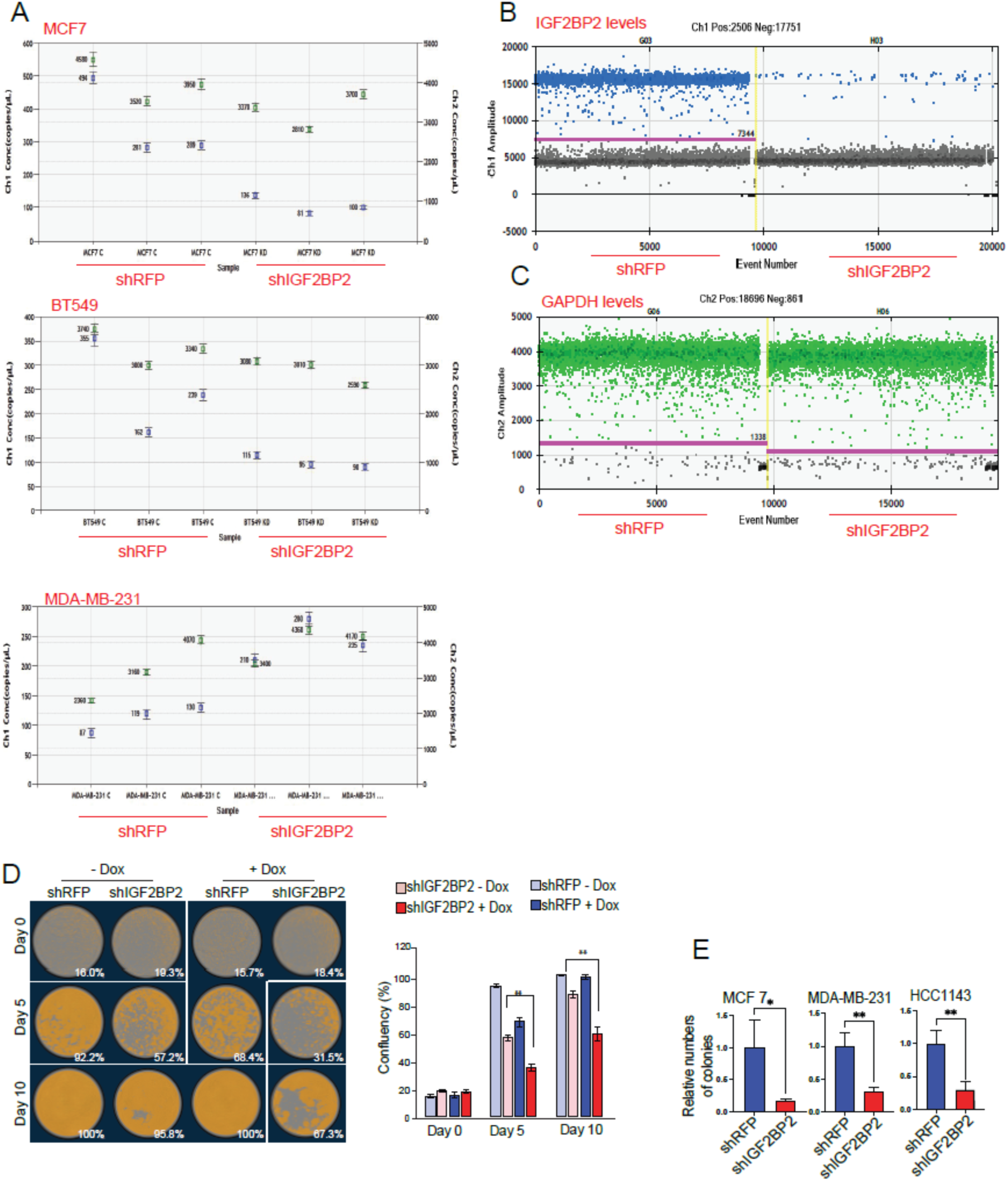
**A.** Absolute quantification (concentration in copies/µL) of GAPDH (VIC-labeled, green) and PLK1 (FAM-labeled, blue) in 3 cell lines, the shRFP control and shIGF2BP2 knockdown samples. **B.** Representative 1D plot showing positive (blue) and negative (gray) droplets based on the IGF2BP2 probe (FAM-labeled) in shRFP control and shIGF2BP2 knockdown cell line samples. **C.** Representative 1D plot showing positive (green) and negative (gray) droplets based on the use of a GAPDH probe (VIC-labeled) in shRFP control and shIGF2BP2 knockdown cell line samples. **D.** Sample images with cell masking are shown in orange, and PLK1-inducible HCT116 cells were quantified using S3-IncuCyte® following the knockdown of IGF2BP2. E. Colony formation assay was performed with the indicated breast cancer cells following knockdown of IGF2BP2. shRFP was used as a control. Colonies were quantified using ImageJ software to determine the number of colonies.

**Table S1.** List of SDL hits identified from the genome-wide shRNA screen with significant weighted differential cumulative changes in PLK-induced versus PLK1-uninduced cells.

**Table S2.** Comparison of all 960 hits with previously published screens associated with mitosis and cell cycle progression.

**Table S3.** List of genes with validated scores based on the work of Marcotte *et al*. or Project Achilles screens used to prioritize PLK1-SDL interactions. Significant p values indicate that PLK1 interacts with the SDL.

**Table S4.** List of genes prioritized using clinical data to capture “naturally occurring” SDL interactions in patients across multiple cancer types.

**Table S5.** List of genes prioritized from 1) ToppGene prioritization, 2) localized to the plasma membrane, and/or 3) located on the cytoband 19p13.2-3.

**Table S6.** List of SDL hits validated by an *in vitro* CRISPR screening approach.

**Table S7.** List of SDL hits validated by an *in vivo* CRISPR screening approach.

**Table S8.** List of sgRNAs used as positive and negative controls in Perturb-seq screening

**Table S9.** List of primer sequences used for genomic cleavage detection.

**Table S10.** Output from mass spectrometric experiments.

## Key resources table

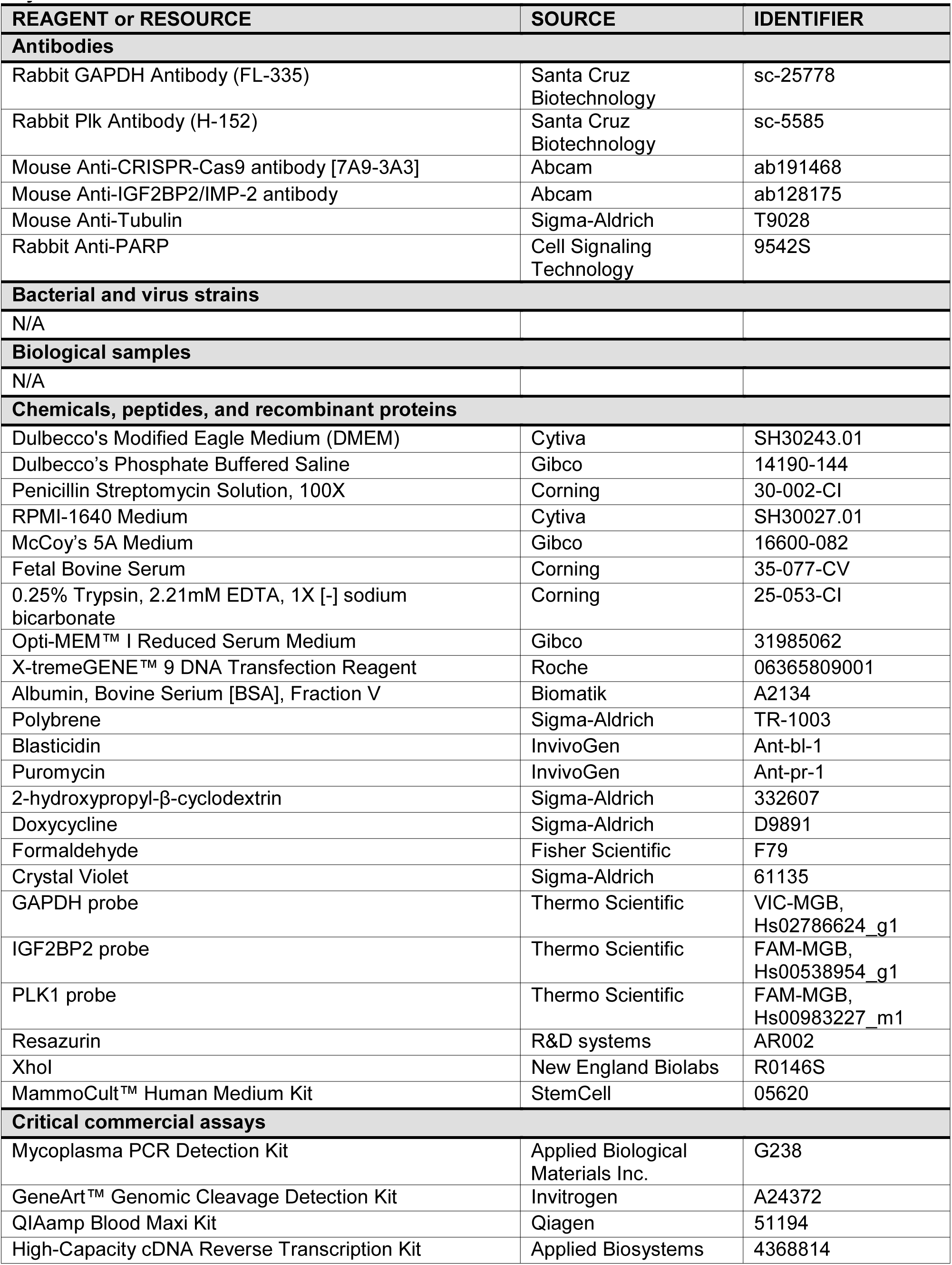
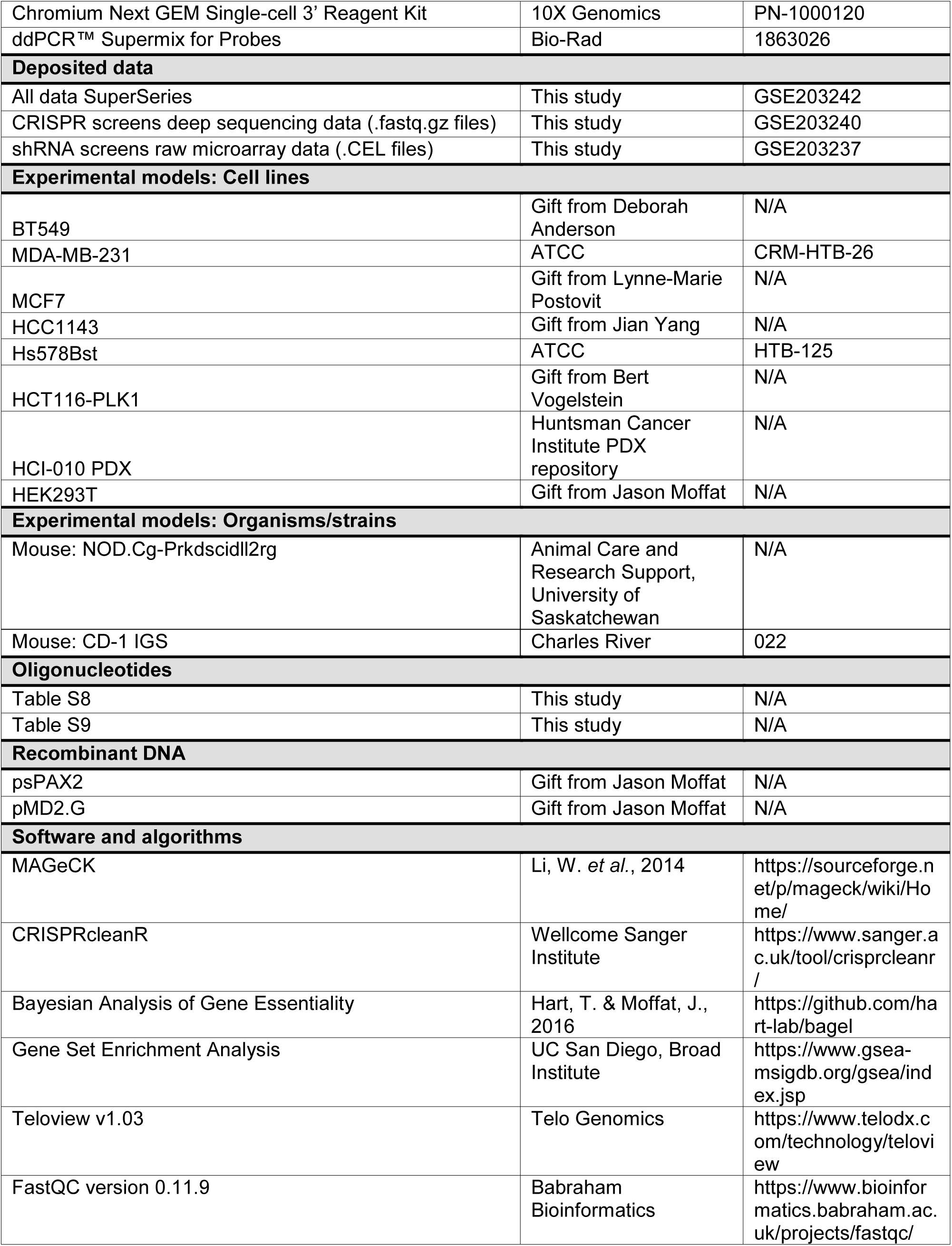
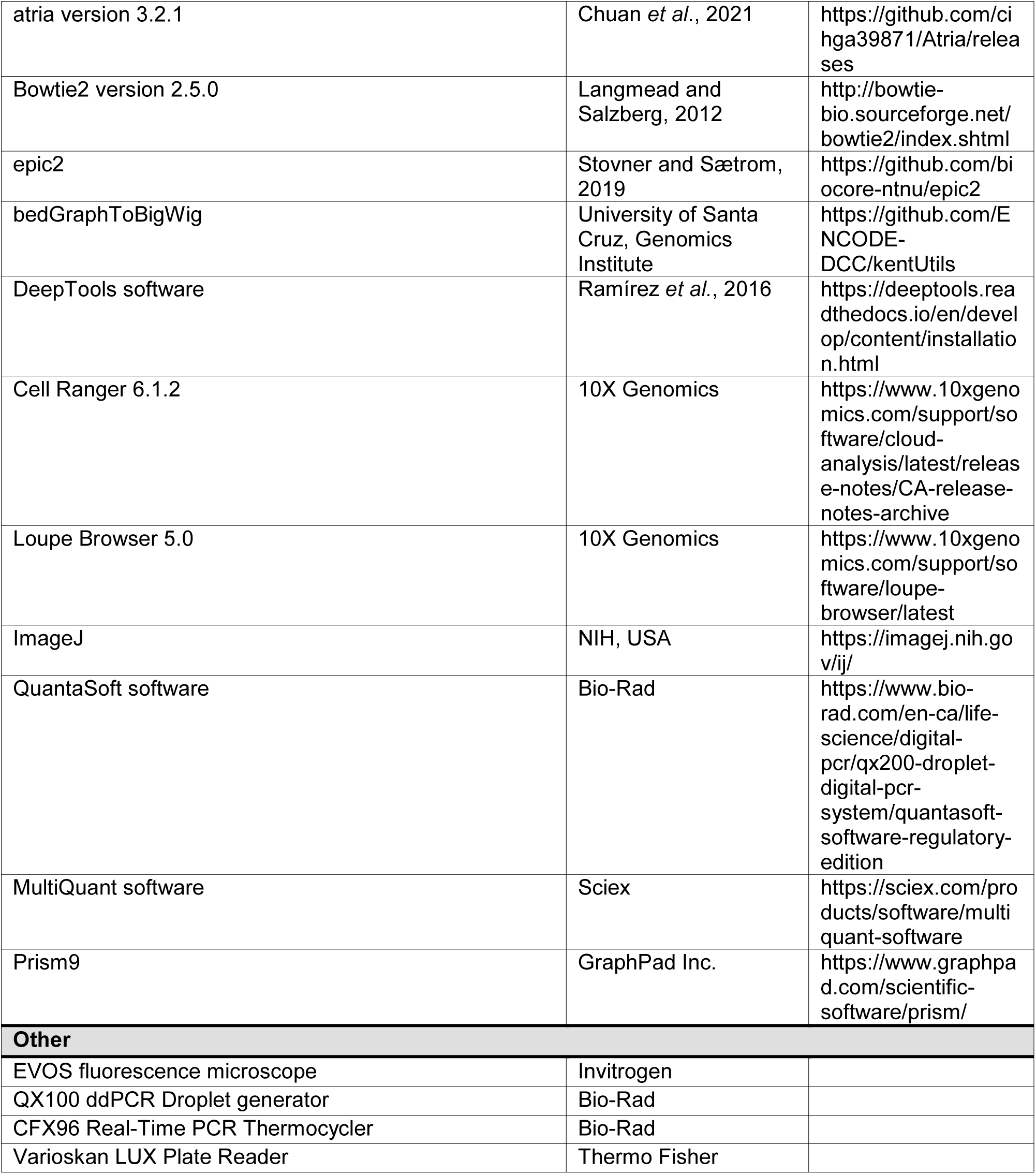

